# AI-derived Protein Structures Validation: AlphaFold2 Models in the Twilight Zone

**DOI:** 10.64898/2026.05.12.724499

**Authors:** Peter Griffin, Giuseppe Deganutti, Kamiya Jadeja, Camila Idigbe, Ludovico Pipito, Lorena Mejuto, Choa Ping Ng, Saffron M. Peck, Jennifer Greaves, Christopher A Reynolds

## Abstract

In any field, unquestioningly accepting artificial intelligence (AI) results should be considered bad practise. Here, we devised a comparative modelling-based strategy for validating protein structures that exploits the well-known observation that protein folds are far more conserved than protein sequences. We identify proteins with a similar fold to the AlphaFold-generated query protein and determine their structural alignment to the query. The hypothesis is that if the sequence alignment coincides with the structural alignment, then the structure is validated. The strategy is implemented on a helix-by-helix and strand-by-strand basis using a multi-template pairwise local profile alignment method that works well into the twilight zone. The method is illustrated by application to the transmembrane transporter PEPT1, for which the structure is known, and the S-deacylases ABHD13 and ABHD16A, for which only AI-generated models exist. ABHD16A is particularly challenging because a sequence alignment search with BLASTp does not reveal any structural homologues and therefore requires work with extremely remote homologues; however, both models are validated through this strategy and are stable during classical molecular dynamics simulations. The ability of the strategy to identify errors is assessed with reference to misaligned ABHD13 models and misfolded decoy proteins.

## Introduction

The ease with which well-trained artificial intelligence (AI) methods can generate accurate results, as in the protein-folding field^1,2^, offers an existential threat to rigorous scientific endeavor as it minimizes the need for professional scientific investigation; consequently, there is a need to validate AI results to ensure they are used with full confidence and understanding. Ever since Anfinsen’s observation that the structure of a protein was encoded in its protein sequence^3^, there has been a huge effort to predict protein structures from sequence alone, e.g. as logged by the numerous CASP competitions^4^. The solution to this 50-year-old protein folding problem has now been largely solved using AI^1,2^, culminating in the 2024 award of the Nobel Prize for Chemistry^5-7^. While the successful AlphaFold and related methods^8-10^ build on sound principles such as multiple sequence alignment, comparative modelling and analysis of correlated mutations, reliance on AI is both a strength, because it usually produces excellent results^11^, and a concern, because, like all AI-based methods, there is little information offered on how the answer, in this case structure, was determined. It is therefore imperative that AI-based results are not used unquestioningly, but rather are interrogated for reliability – not just in protein folding or other areas of science but in all areas of life.

Our validation method draws heavily on the principles of comparative modelling^12,13^, which requires (i) the identification of one or more structural homologues (templates) for the unknown protein query, typically identified using a BLASTp^14^ search and (ii) a multiple sequence alignment of the target and templates, typically generated using sequence alignment software such as Clustal Omega^15^. The generation of the target structure from the templates is critically dependent on the quality of the sequence alignment, but comparative modelling struggles when the percentage identity (PID) is below 30%^16-18^, particularly so in the case of extremely remote homologues (< 20% identity)^19^.

The approach we propose effectively reverses this process. Since there is an AI-derived query structure, structural homologues for use as validation templates are found by searching in structural space rather than sequence space using the Dali protein comparison webserver, which is more effective than BLASTp-based sequence searches. This is because structure is better conserved than sequence, as shown for example by the Homstrad^20^, Dali^21,22^, SCOP^23^ and CATH^24-26^ databases. Using the principles of comparative modelling, it follows that if the AI-generated structure is correct, then the sequence alignment of the query and template should agree with that generated by the Dali structural alignment. The biggest problem in this strategy is the alignment between low-PID sequences, as stated above. However, we have developed a sequence alignment method designed to work^27-29^ in or below the twilight zone (20-35%)^30^, enabling the use of remote structural homologues in the validation process. Our method was first used to develop a model of Androglobin validated by experiment^29^, when previous approaches, based on incorrect alignment^31^, stated the protein could not be expressed^32^; remarkably, the mean PID of Androglobin to other globin templates of known structure was around 14-18%^27-30^.

Here, we further validate our method on two important drug targets: PEPT1, a dipeptide transporter of known structure involved in drug transport^33^; and ABHD16A, an important cerebellar phosphatidylserine lipase with neuroinflammatory implications^34^. We have also applied the method to ABHD13, a previously uncharacterized protein that, together with ABHD16A, reversed S-acylation, a widespread post-translational lipid modification of proteins that is particularly prevalent in the brain and nervous system^35,36^. We have recently reported, exploiting AlphaFold models, that ABHD13 and ABHD16A selectively deacylate the SNAP25 family of Soluble-N-ethylmaleimide-sensitive factor attachment protein receptors (SNARE) proteins responsible for vesicular membrane fusion (see the accompanying preprint Mejuto et al., bioRxiv, 2026, doi: 10.64898/2026.01.21.700842), underscoring the relevance of this validation strategy.

## Methods

### Template Identification using BLASTp

The amino acid sequences for the query AlphaFold2 proteins were submitted to BLASTp (https://blast.ncbi.nlm.nih.gov/Blast.cgi?PAGE=Proteins) to identify homologues from the Protein Data Bank.

### Template Identification using Dali

The PEPT1 protein structure^33^ used was the open occluded structure, 7pmw, as it has the smallest RMSD to the other PEPT1 structures (7pmx, 7pmy, 7s8u, 7ps1). The AlphaFold2 models of human ABHD13 and ABHD16A (Figure 1A,B) were downloaded from the AlphaFold2 Protein structure database (https://alphafold.ebi.ac.uk/entries%20Q7L211%20and%20O95870, accessed on 17 November 2022) and submitted to the Dali protein comparison webserver (http://ekhidna2.biocenter.helsinki.fi/dali/); the protein hits were identified, and the associated structural alignments downloaded. The structural alignments of each helix and strand to the query protein were scored using the BLOSUM 62 matrix, and the scores were summed to give a score for each template. For ABHD13, seven high-scoring templates were identified as suitable for the validation process. For ABHD16A, because the protein hits were more remote homologues, 5 templates were chosen for each helix or strand (56 in total), subject to the constraint that the template has the correct secondary structure and positive alignment scores

**Figure 1.**
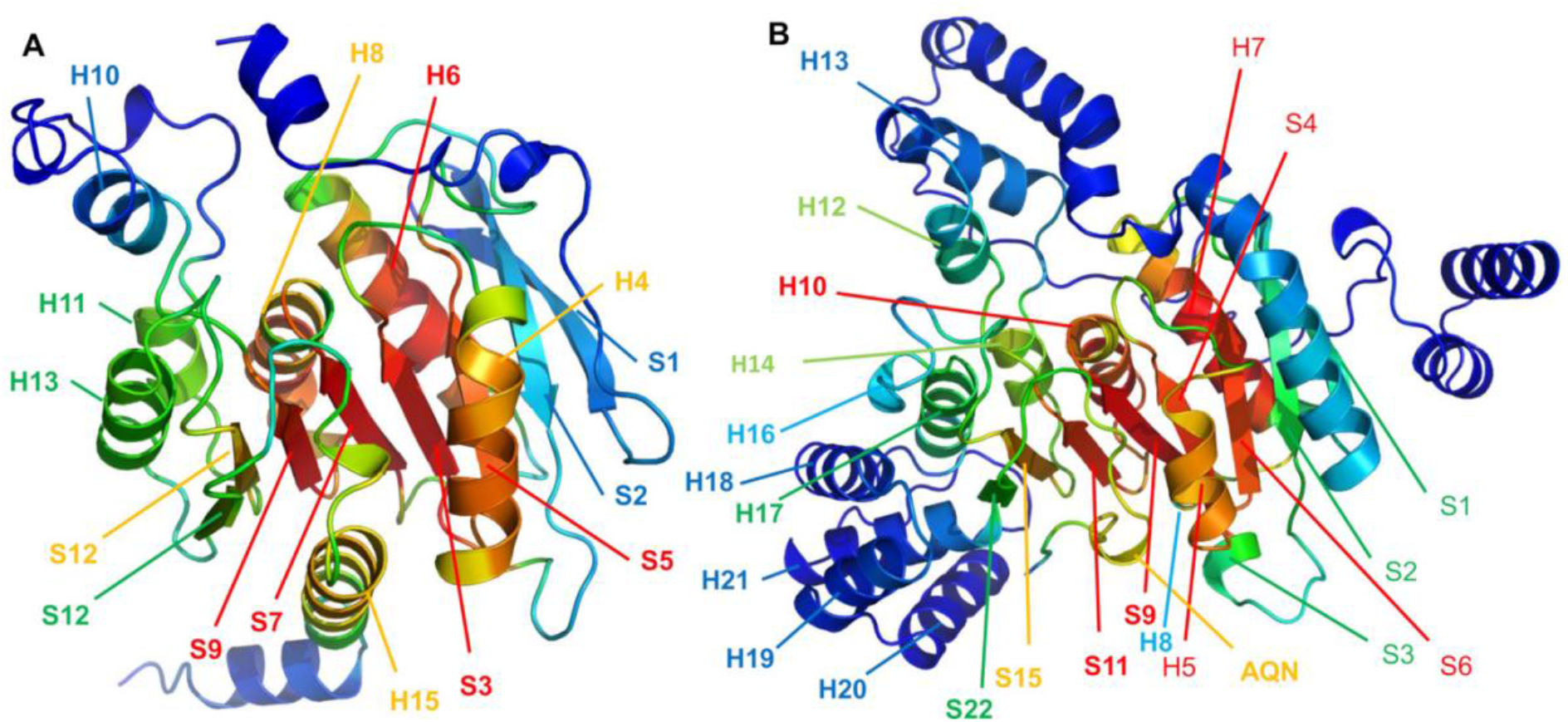
Template structural Landscape for (A) ABHD13 and (B) ABHD16A. The number of structures that align to ABHD13 or ABHD16A are colour coded (blue low, red high). The assessed structural elements are labelled. Some helices and strands are not labelled because there was no comparable structure in the Dali database. The N-terminal membrane binding regions were omitted because no hits were identified by Dali.

### Multi-template pairwise local sequence alignments

For each of the secondary structural elements, the alignment was determined using in-house multi-template pairwise local alignment (MTPLA) code, which was designed to work in situations of low PID^28,29^. The essence of the method is illustrated with reference to Figure S1.

Figure S1A shows the lead (human ABHD13) sequence for the 65 homologous sequences for ABHD13 and the lead (5g59, a bacterial esterase) sequence for the 111 homologous sequences for protein 5g59, along with the secondary structure for ABHD13. For the 7 pairs of sequences (1 for each of the templates), the alignment is scored in the alignment window using the BLOSUM62 matrix. The alignment is then adjusted by moving the bottom sequence (ABHD13) one position to the left (the −1 alignment, see Figure S1B) and scored. The process is repeated for all positions through −1 to −8 (e.g. −3, as in Figure S1C) and for all positions from 1 to +8 (e.g., +3 in Figure S1D). The highest scoring position gets one vote (e.g., Figure S1A); the other positions get zero votes (Figure S1B,C). The process is repeated for all sequence pairs (7215 in the case of ABHD13 / 5g59), and the position receiving the most votes is noted. The same procedure is then applied by moving the top sequence (in this case 5g59) in the same way (Figure S1D,E). Note that when 5g59 is moved 3 places to the right, the alignment corresponds to that obtained by moving ABHD13 3 places to the left (Figure S1C, Figure S1E). Votes are received in the same way, and the total score is obtained as the mean of the score for moving the lower sequences and that obtained by moving the upper sequences. However, while the alignments in Figures S1C and S1E are identical, in these two cases, the local alignments scored are different because the residues within the scoring box differ; the size of the scoring box generally equates with the length of the helix or sheet but may be shorter if there are too few usable templates. For this reason, the most reliable score is obtained for the 0 alignment, so in general, if the results indicate +3 is the correct alignment, then the lower sequence should be adjusted leftwards by three positions so that 0 is the correct alignment and the alignment re-determined.

This procedure is repeated for all seven templates, noting that where there are gaps in the alignment (e.g. helix 4 of 4×00, helix 10 of 2i3d) or there is a change in the secondary structure (e.g. helix 10 of 5hdf), some secondary structural elements may have fewer than seven templates. The results for all valid templates are scaled between 0 and 1 and multiplied together; this has the effect of re-enforcing strong signals and decreasing noise. The method does not require the strongest signal to be at 0 as long as the 0 alignment has a reasonable score, e.g. above 0.25. The truly random score is 0.06 (1/17) for a window of ±8 and 0.04 (1/25) for a window of ±12; a window of 12 was used for most helices and strands, but only 8 is displayed.

### Simulations

The simulation methodology is as reported in the accompanying preprint (Mejuto et al., bioRxiv, 2026, doi: 10.64898/2026.01.21.700842).

## Results

### Homologues identified by BLASTp

The structures identified from the Protein Data Bank for PEPT1^33^ were 2xut, a PEPT1 homologue from *Shewanella oneidensis*, and 4ikv and 6ei3, both bacterial POT family transporters. The results of a BLASTp search for the human sequence of ABHD13 (Uniprot code Q7L211) are given in Table S1; the corresponding search for the human sequence of ABHD16A (Uniprot code O95780) yielded no hits. This indicates that the generation and validation of the AlphaFold model of ABHD16A is considerably more challenging than for ABHD13. Even for ABHD13, the overall percentage identity (average 27%) of the hits in Table S1 lies within the twilight zone where sequence alignment is difficult, and the average coverage is only 52%.

Submission of the ABHD13 and ABHD16A models to the Dali protein comparison webserver yielded 999 and 1477 structural alignments, respectively. However, the number of close structural alignments for each region of the sequence, determined by residues with capital letters in the structural alignment, varied between 0 and 329 for ABHD13 and between 0 and 126 for ABHD16A. Consequently, for ABHD16A, the number of available templates varied for each helix or strand.

### Template identification

The ABHD13 templates identified were 2i3d (alpha-beta hydrolase of unknown function), 2o2g (Dienelactone hydrolase), 3pf8 (cinnamoyl esterase), 4ao8 (cold-adapted esterase), 4×00 (putative aryl esterase), 5g59 (Pf2001 esterase) and 5hdf (Streptonigrin methylesterase A). The sequence alignments derived from the structural alignments for ABHD13 and ABHD16A are shown in Figures 2, 3, and Figures S2 and S3, respectively, along with the chosen template sequences for each helix or strand.

**Figure 2.**
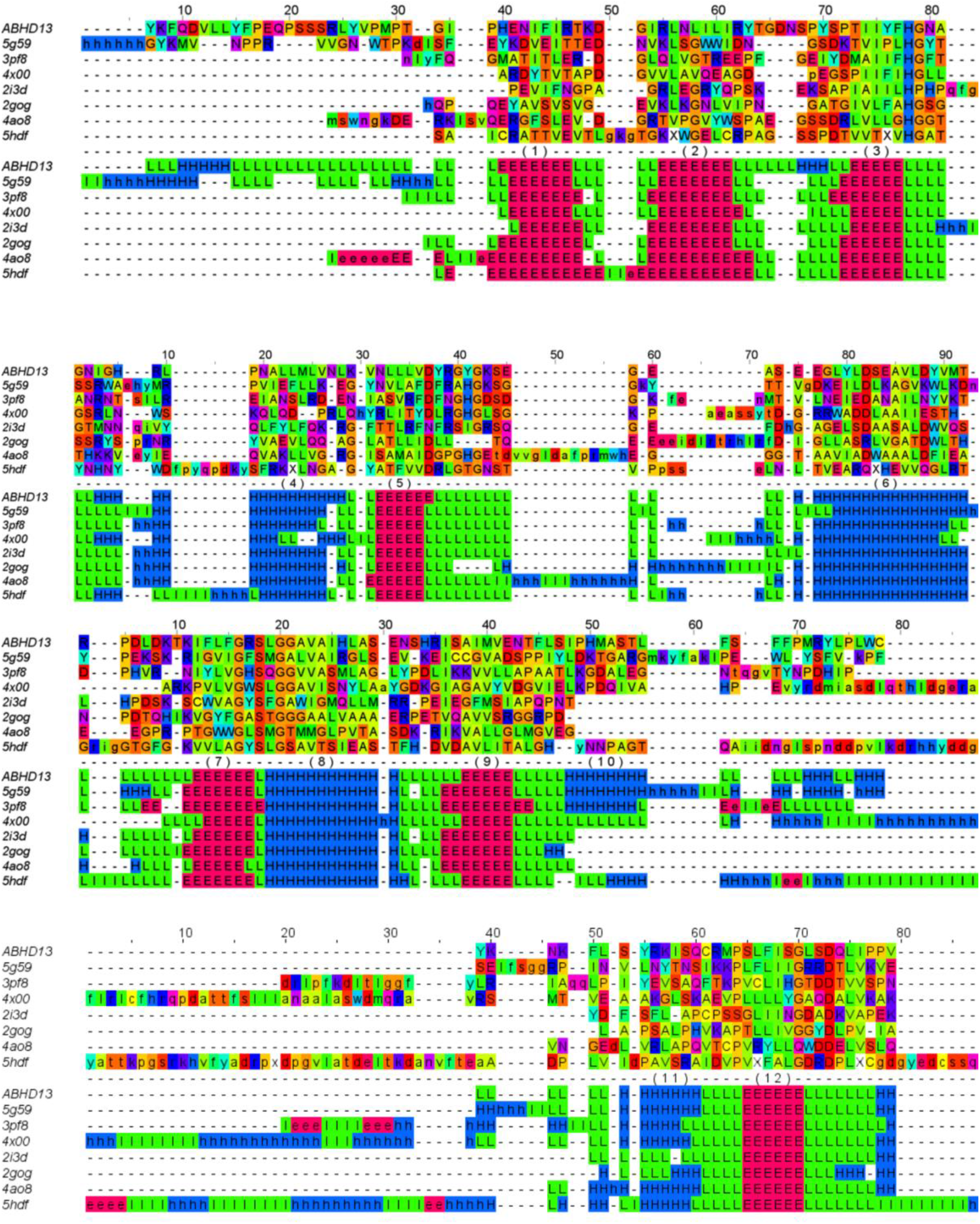
The structural alignment between ABHD13 and the 7 templates, as determined by the Dali protein structure comparison server. The 7 templates were chosen because of a relatively high similarity score over the helices and sheets, as determined using the BLOSUM62 similarity matrix.

**Figure 3.**
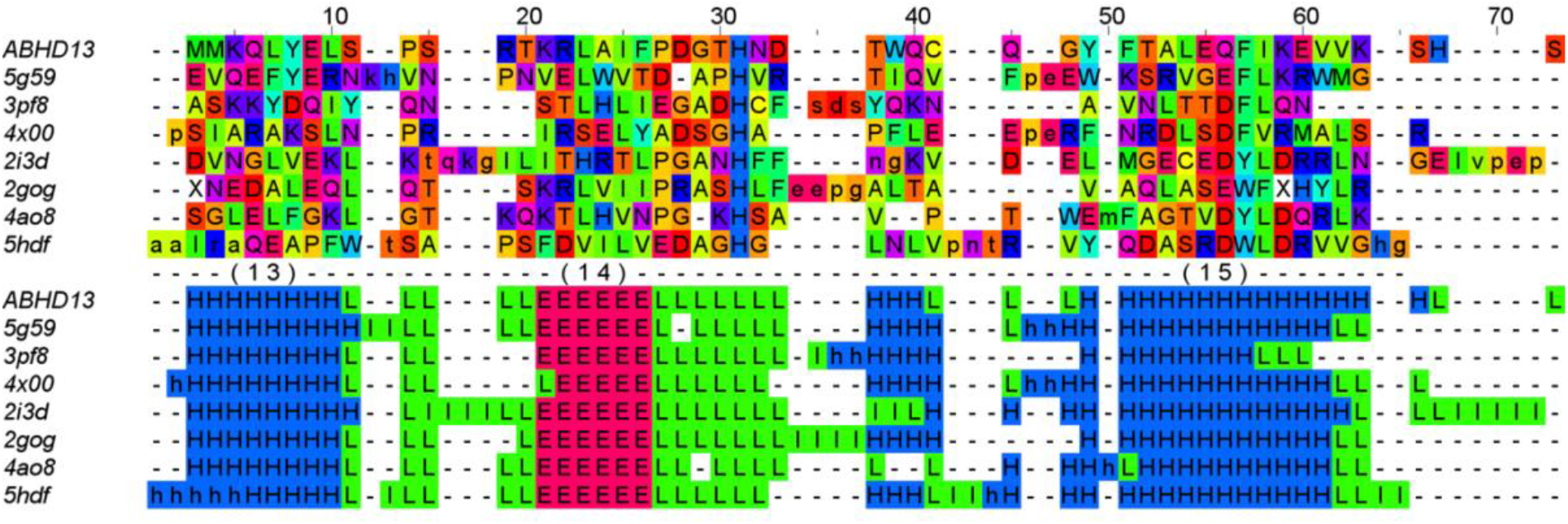
The structural alignment between ABHD13 and the 7 templates (continued), as determined by the Dali protein structure comparison server. The 7 templates were chosen because of a relatively high similarity score over the helices and sheets, as determined using the BLOSUM62 similarity matrix.

### PEPT1 validation

The 7pmw PEPT1 cryo-electron microscopy structure^33^ was used as a control to ‘validate’ the cryo-electron microscopy structures. The alignment results for the 12 transmembrane helices are shown in Figures 4 and S4. For each transmembrane helix, the highest peak is at 0; this validates the method for this case. For some helices, e.g. helices 1, 3 and 4, the percentage identity is high (36.7 - 63.2), but all other helices have one or two templates, where the alignment is in the twilight zone. For helix 12, the percentage identities to the templates are 10.1%, 19.7% and 17.5%; these are well within the extreme remote homology zone, and so helix 12 represents a particularly rigorous test of the method. Given that the method works for validating a known experimental structure, we applied it to our target AlphaFold structures.

**Figure 4.**
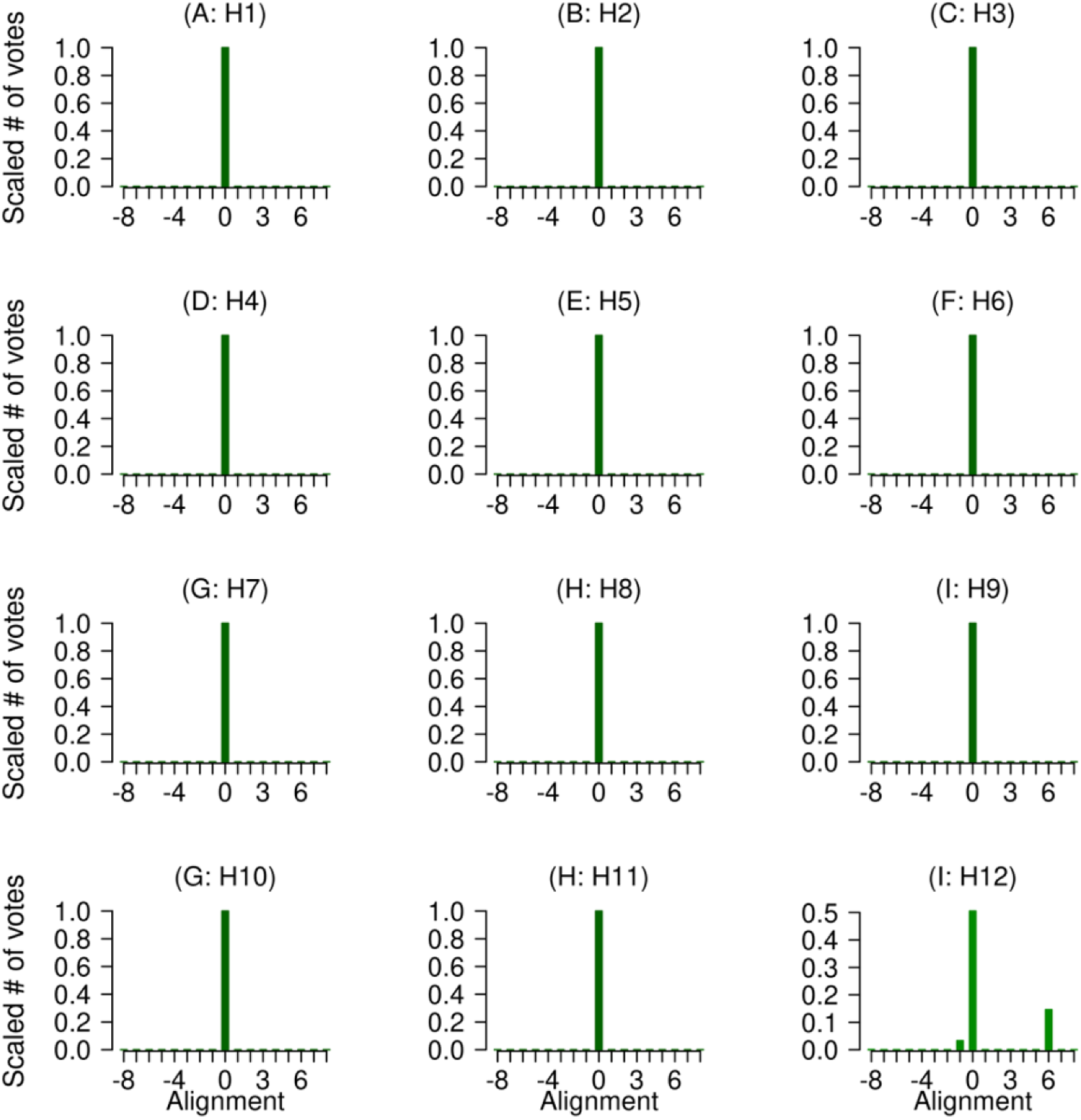
‘Validation’ of the PEPT1 cryo-EM structures, using the structural alignment of the 2xut, 4ikv and 6ei3 bacterial templates. The local multiple-template alignment method validates the structural alignment since the highest score is at 0 for each of the 12 transmembrane helices. Level 1 validation is shown in dark green (e.g. H1), level 2 in a less dark green (e.g. H12).

### ABHD13 and ABHD16A validation

The interpretation of the results is discussed in detail for ABHD13 (Figures 5 and 6); similar principles apply to ABHD16A (Figure 6, Figure S5 and S6). For ABHD13 strands 2, 7, 9-10, 12 and 14 and helix 8, there is a clear bar of height 1.0 (or close to 1.0) at 0 on the alignment axis, with no other bars of comparable height. The corresponding secondary structural elements for ABHD16A with a clear bar of height 1.0 (or close to 1.0) are strands 1, 3, 6, 9, 12 and helices 5, 8 and 10. A bar of height 1.0 (or close to 1.0) gives the strongest validation criteria (i.e. level 1) of these particular helices or strands within the AlphaFold model (see Table S2 for validation levels). A clear example of this is seen for strand 12 (SLFISG) in Figure 5, where the peak is 1.0, indicating that the individual alignments of ABHD13 against each of the seven templates also gave an alignment of 0, as shown in Figure 6 (Figure 5 is derived from Figure 6 by multiplying the 7 sets of peak heights together). The two weak peaks at alignment 1 for 2o2g and 4ao8 (Figure 6) disappear in Figure 5 due to the process of scaling the peak between 0 and 1 and multiplying, since the product contains multiple small numbers.

**Figure 5.**
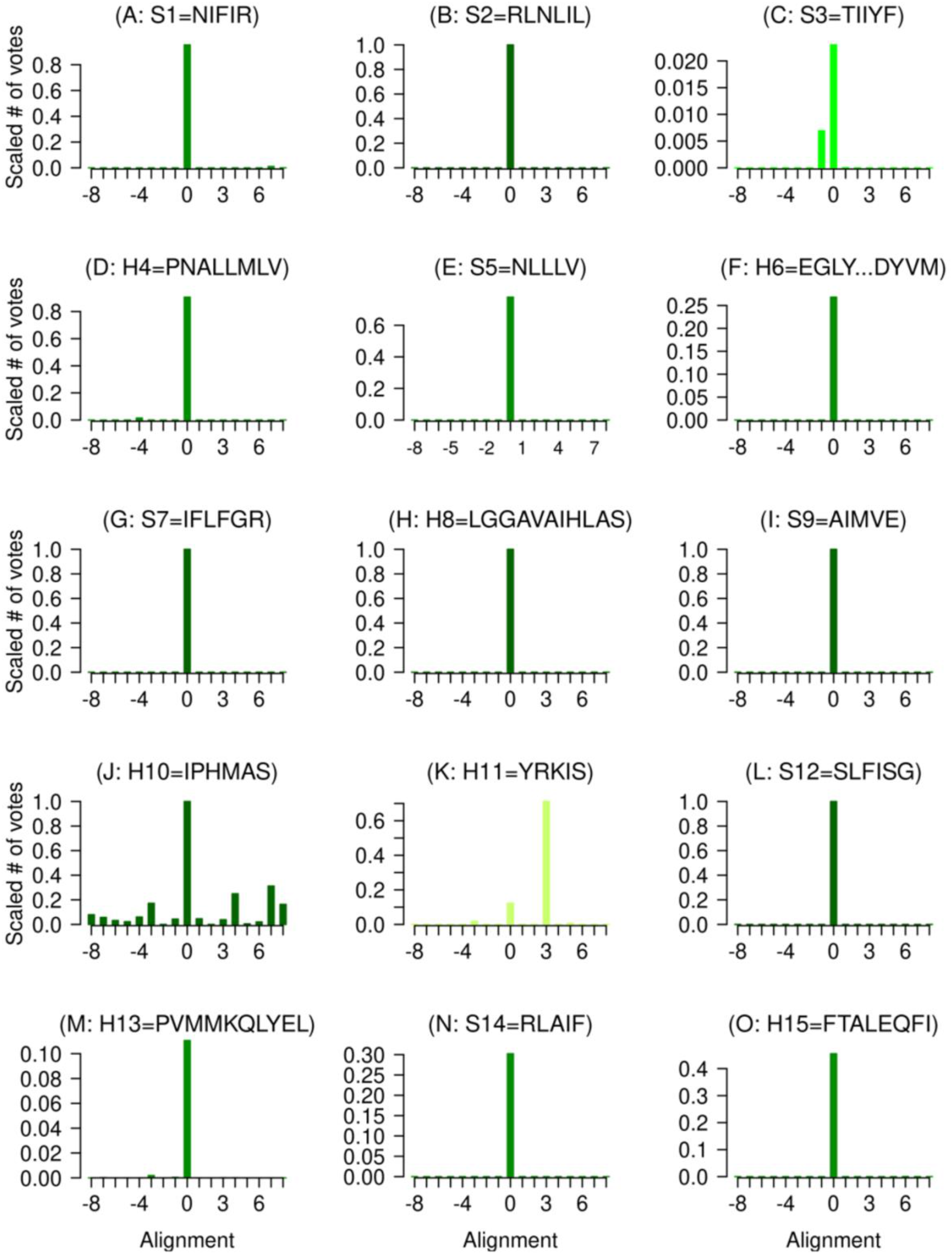
Validation of the AlphaFold2 ABHD13 structure, as determined by the multiple template alignment method over the 8 sheet and 7 helical regions. Level 1 validation is shown in dark green (e.g. S2), level 2 in a less dark green (e.g. S1), level 3 in green (e.g. S3) and level 4 validation in dark olive green (e.g. H11).

**Figure 6.**
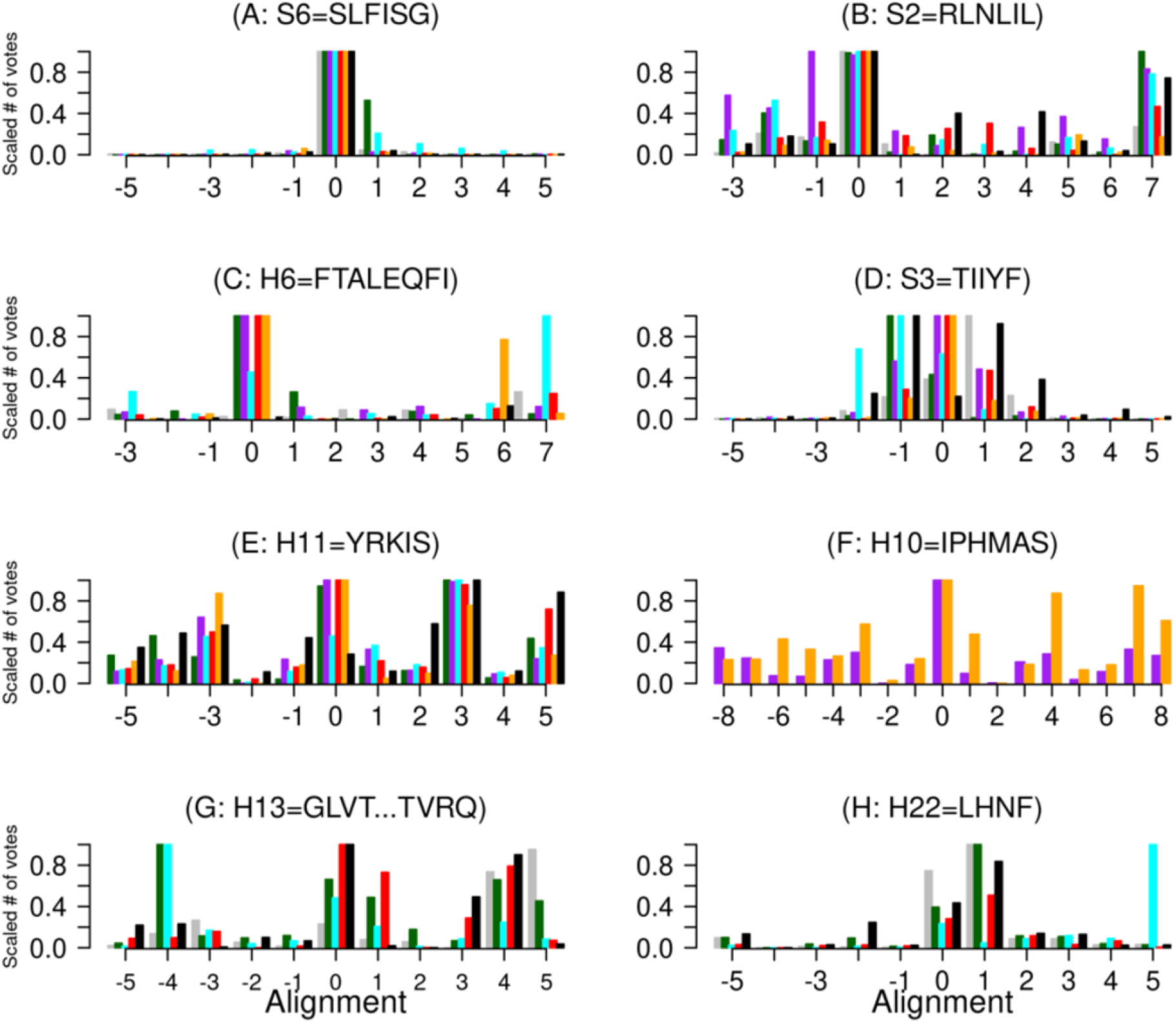
Analysis of individual templates in the validation of the AlphaFold2 ABHD13 and ABHD16A structures, as determined by the multiple template alignment method over the 8 sheet and 7 helical regions. ABHD13 Colour code: grey is 2i3d, dark green is 2gog, purple is 3pf8, cyan is 4ao8, red is 4×00, orange is 5g59, black is 5hdf. A-F: ABHD13; G-H ABHD16A. G colour code: 2a3b is grey, 4×4w is dark green, 5w66 is cyan, 5yu7 is red, and 4j8s is black. H colour code: 3o4j is grey, 6wym is dark green, 3fnb is cyan, 7dwc is red, and 1u2e is black.

For strand 2 (RLNLIL), there is a strong peak of almost 1 at 0 (Figure 5B), but analysis of the seven individual components in Figure 6 shows a more complex scenario, as there are individual peaks of 1 at alignments −1 (3pf8), and 0 and 7 (2o2g). However, the peaks for other templates at −1 and 7 are small and so disappear during multiplication, leaving only the peak at 0, supporting the modelling solution of this structural element by AlphaFold.

For helix 15 (FTALEQFI) (Figure 5O and Figure 6C), there is a single peak in Figure 5M, but this is reduced because 3pf8 gives an alternative main peak at alignment 7. However, 3pf8 also has a peak of 0.46 at alignment 0, and this contributes to AlphaFold validation. Thus, Figure 6 shows that for this multi-template alignment method to work, all the templates do not need to give the highest score to the zero alignment, as long as the zero alignment does not receive a low score. This criterion of a strong peak (not necessarily as large as 1.0) at zero gives the second strongest degree of validation (level 2). Level 2 validation is seen for strands 2 and 11 and helices 3, 5, 7, 14, 16, 19, 21 and 23 in ABHD16A (Figures S5, S6). Helix 15 lies at the C-terminal region of the templates, and there is insufficient sequence data in the multiple sequence alignments to generate local alignments for the 3pf8 or 5g59 templates; hence, the use of only 5 templates.

Strand 3 (TIIYF) has a strong peak at 0, with a smaller peak at −1 (Figure 5C). The data for the associated individual proteins shown in Figure 6D shows strong peaks of 1.0 at both the 0 and the −1 alignments. This data provides strong validation of the AlphaFold model, partly because the alternative peak in Figure 5C is relatively weak, and partly because the 0 alignment is well-represented by strong peaks in Figure 6D. This criterion, where the peak at 0 is the strongest, but there are smaller competing peaks, is the third level of validation (level 3, Table S2). In ABHD16A, level 3 validation is seen for helices 12, 17, 18, 19, 20 and 23; helix 3 could be assigned to this level, but the peak at 1 is quite small. Helix 19 falls into this category because the PID to the templates is low; helix 20 falls into this category because there are only two templates, which limits the filtering effect of multiplying by small numbers.

Elsewhere, namely helix 11 in Figure 5K and helices 13 and 22 in ABHD16A (Figure S4M and Figure S5H, we see a similar situation in reverse. For ABHD13, the individual protein data, shown in Figure 6E shows support for both the 0 alignment (2o2g, 3pf8, 4×00 and 5g59) and alignment +3 (2o2g, 3pf8, 4ao8, 4×00 and 5hdf). The stronger peak at alignment 3 (Figure 5K) could be taken to suggest that the AlphaFold model is in error in helix 11, but we conclude that the peak at 0 shows that the AlphaFold helix 11 structure is *compatible* with the data, and so the structure is validated. This is the fourth level of validation (Table S2). Note that 2o2g, 3pf8, 4×00 score strongly at two different alignments – this is a common theme. Figure 6E also shows peaks at −3 and 5, but these are much weaker and are almost reduced to zero by the multiplication process. Figures 5K and 6E are based on just 6 proteins since the overall Dali-derived structural alignment shows a gap in helix 11 of 2i3d. The mean percentage identity between ABHD13 and the templates is 8.6% for helix 11, which is low (midnight zone), so results for this region need to be interpreted with caution. Likewise, Figure 6G,H shows that there is support for the 0 alignment in the individual alignments for both H13 and H22 of ABHD16A, again showing that the AlphaFold structure is *compatible* with the data for these helices; in each case. The mean PID between query and templates is 10% in both of these cases. Figure S3 shows that the ABHD16A helix 13 region is not fully helical for four of the common templates. Helix 13 is, however, helical for the templates used, which are specified in Table S3.

There are gaps in the 4×00 and 5g59 helix 13 template multiple sequence alignments, and so Figure 5M is based on just 5 templates. Helix 10 is likewise based on just 2 templates; consequently, there are multiple small side peaks as the filtering effect of multiplication is reduced. Both 3pf8 and 5g59 gave the main peak at 0, as shown in Figure 6F. For strand 14, which is clearly validated (Figure 5N), the percentage identity between ABHD13 and the templates is 6.5%, which again is well within the midnight zone (percentage identity < 10%).

Overall, the set of results in Figures 5 and S5, and S6, is quite remarkable as the mean percentage identity (PID) between the templates and the ABHD13 model is 17.4% for ABHD13 (Table S4) and 16.1% for ABHD16A (Table S3), which is well within the twilight zone. The general conclusion is that the AlphaFold model has been validated by the multi-template pairwise local alignment model, and so it can be used in structural studies with confidence. Nevertheless, the 4 different levels of confidence reported suggest regions where additional examination may be beneficial, e.g. by molecular dynamics.

### Misaligned protein models

The ability of our validation strategy for detecting misalignment within a protein model has been tested by introducing random misalignment errors (between ±4) into the ABHD13 structural alignment. The results, Figure S7, show that in general, the peak shifts from 0, so as to indicate the extent of the alignment error. Thus, in Figure S7A, where an alignment error of 4 was introduced (ABHD13 motif moved 4 positions to the right), the peak is at −4, indicating that the query ABHD13 sequence needs to be moved 4 positions to the left to gain the correct alignment. The only ambiguous result is in Figure S7D, where an alignment error of −1 gives rise to almost equivalent peaks at 0 and 1. However, the equivalent peak at 0 in Figure 5D is unambiguous. The discrepancy between Figure 5D and Figure S7D arises because the alignment boxes corresponding to 0 and 1 in Figures 5D and S7D are different. As described in the methods, for optimal validation, the alignment should always be adjusted to ensure that the correct alignment occurs at 0.

### Misfolded protein models

Six misfolding decoys were taken from the Decoys ‘R’ Us database of incorrect conformations^37^: the sequences of proteins 2cro and 1sn3 mapped onto the structure of 2ci2, the sequences of 2ci2 and 1sn3 mapped onto the structure of 2cro, and the sequences of proteins 2ci2 and 2cro mapped onto the structure of 1sn3. In an attempt to validate the decoys, which are misfolded but designed to have native-like features, we assessed the structural alignments shown in Figure S8, as shown in Figure S9. Thus, Figure S9A shows the alignment of the 2cro sequence against 2ci2 over the region of helix 1 of 2ci2 (orange) and the alignment of the 1sn3 sequence against 2ci2 over the region of helix 1 of 2ci2 (green). On the (false) assumption that the 2cro and 1sn3 decoys have the same fold as 2ci2, it would be reasonable to scale and multiply the alignment scores to generate a product (black bars in Figure S9). Several observations arise that are consistent with the decoys having the incorrect fold. Firstly, 2cro gives rise to multiple high peaks in different positions: this is not so apparent for 1sn3 in Figure S9A, but it is observed in Figure S9B and elsewhere in Figure S9. Examples of this are shown in Figure 6 for individual (rather than product) alignment, but it is more marked in Figure S9. Secondly, while some individual high peaks occur randomly at 0, generally, the high peaks are elsewhere. Thirdly, while in a small number of cases, multiplication reinforces the evidence for certain alignments, the peak is not at zero. Moreover, multiplication generally minimizes the signal, as would be expected for misaligned structures or misalignment. The percentage identity at the zero alignment is in the range of 4±4, which is to be expected for random sequences.

### The effect of PID on alignment scores

The distribution of secondary structural elements according to percentage identity for PEPT1, ABHD13, and ABHD16 is shown in Figure 7A. Almost 84% of the data lies within the twilight zone (<35%), 59% of the data lies within the zone of extreme remote homologues (<20%) and 24% lies in the midnight zone (<10%), and so this data set clearly presents a challenge for any validation method. Significantly, the dependence of the individual alignment score on percentage identity is shown in Figure 7B (with the proviso that the AlphaFold structures are correct). Seventy five percent of the scores are 1.0, showing that the single template alignment method works in the majority of cases. More significantly, 95% of the scores are above 0.25, meaning that they can strongly contribute to the multi-template method. There is only one score below 0.9 for any alignment with a percentage identity above the extreme remote homologue zone (20%), namely 0.63 at 20.1%. Figure 7C shows the growth in the errors as the percentage identity decreases. Taking scores below 1.0, 0.9 and 0.8 as in error, the error does not rise to 5% until the percentage identity drops to 15%, 11% and 10.4%, respectively. However, with the focus on the multi-template approach, where scores as low as 0.25 can be useful, the error does not rise above 5%. Only two data points in Figure 7B correspond to scores below the truly random value of 0.06 for a window of ±8 or one below 0.04 for the more usual window of ±12; the data points have a percentage identity of 5.8 and 3.9, respectively.

**Figure 7.**
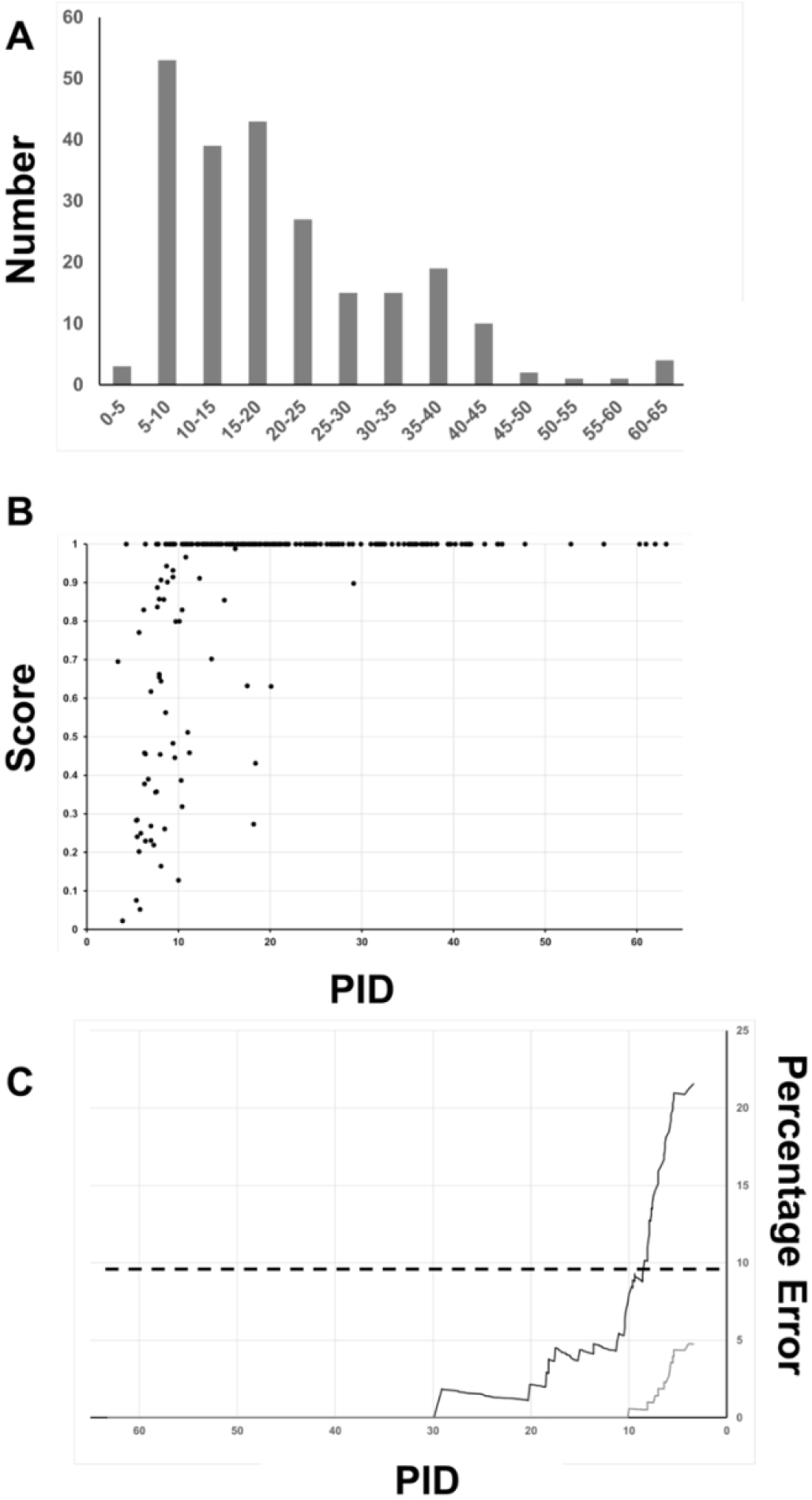
Percentage identity and alignment score in fold validation. A. The distribution of secondary structural elements according to percentage identity. B. The relationship between the single template alignment score and percentage identity. C. The error rate as a percentage of identity is decreased: the top curve relates to scores above 0.9; the bottom curve relates to scores above 0.25.

There is no correlation between the number of available templates and the validation level, and neither is there a correlation between the length or identity of the secondary structural elements and the validation level (Figure 7).

### Molecular Dynamics Simulations

The RMSF of ABHD13 and ABHD16A is shown in Figure 8. Clearly, the extent of motion observed is system-dependent, and large conformational changes would require simulations in the millisecond or second timescale. The large RMSF at the unconstrained C-terminus of ABHD13 is to be expected. There is no clear correlation between the RMSF data and the level of validation, so although helix 11 of ABHD13 (validation level 4) is near a region of higher RMSF, because of the proximity to a long loop, it shows similar RMSF values to strand 2, which has the highest level of validation. Consequently, the results are consistent with those obtained from µs simulations of stable X-ray structures, and so the molecular dynamics simulations indicate that there are no problems with the structures.

**Figure 8.**
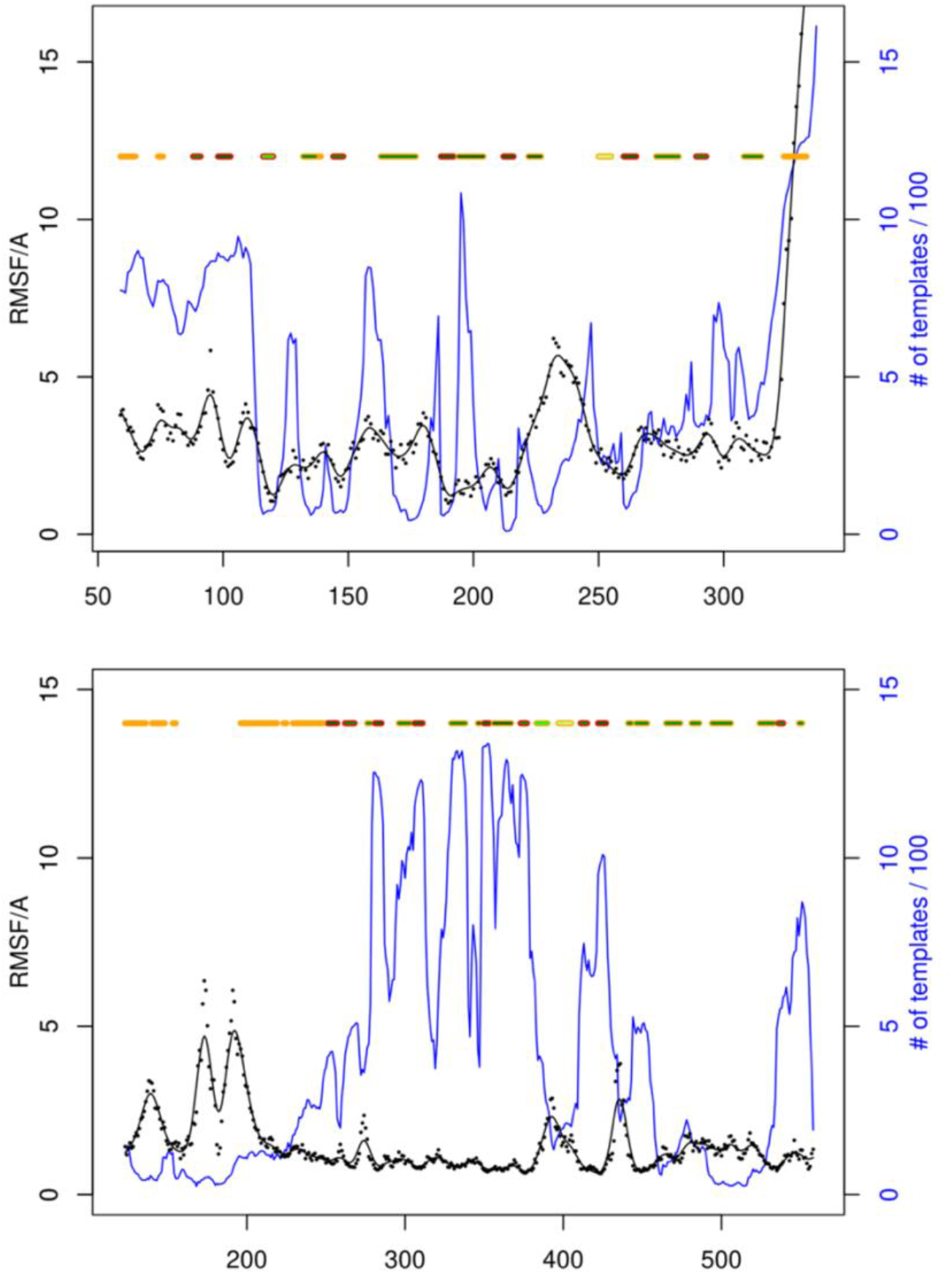
RMSF (black) determined by classical molecular dynamics simulations on (A) ABHD13 and (B) ABHD16A. The number of structurally aligned residues determined by Dali is shown in blue. The motifs are shown above with a red outline for sheets and an orange outline for helices. The shade of green displays the validation level (see Figure 5); solid orange denotes helices where there were no contiguous corresponding motifs in the structural alignment.

## Discussion

The recent advent of experimental quality protein structure^4^ through AI-based computational methods such as AlphaFold^1,2^ heralds an era of greatly expanded opportunities, not least in the field of medical research. However, as in all AI-based methods, there are potential problems if the results are not critically evaluated. Our strategy for evaluating AI-derived protein structures has been tested on the PEPT1 dipeptide transporter, which is important as it can be targeted for drug delivery. It has also been applied to two S-deacylases of potential medical importance, namely ABHD13 and ABHD16A^38,39^. Successful validation of ABHD13 has given confidence in MD simulations carried out in support of experiments showing that ABHD13 and ABHD16A play a key role in deacylating SNAP25 proteins and in explaining how ABHD13 distinguishes between two highly similar *S*-acylated substrates - SNAP25a/SNAP23 and SNAP25b - and between distinct *S*-acylation sites within the same protein.

Our validation strategy inverts the principles of homology modelling and exploits the observation that fold is more conserved than sequence^20-26^. There are many methods for identifying whether proteins have a similar fold^21,38^,40,41; here we have used the Dali protein comparison webserver. The structure is validated if the protein sequence alignment coincides with the protein structure alignment. The huge difficulty with this strategy is that many of the required alignments lie within the twilight zone (35-10% sequence identity) or even into the midnight zone (10-0% sequence identity) where sequence alignment is difficult or verging on impossible. To combat this problem, we have used the multi-template pairwise local profile alignment method, an in-house sequence alignment method shown to give accurate results within the twilight zone^29^. As a local un-gapped sequence alignment method, it is applied to each helix and strand. In the case of PEPT1, the method correctly validated each of the transmembrane helices, including transmembrane helix 12 where the percentage identity lies in the zone of extreme remote homologues^19^.

BLASTp identified eleven homologues of known structure for ABHD13 but for ABHD16A none were found, making the ABHD16A AlphaFold model a more difficult structure to validate. This was alleviated by selecting distinct Dali-identified templates for each secondary structure element rather than for the protein as a whole, as in PEPT1 and ABHD13. The downside is that more multiple sequence alignments are required (56, as opposed to 3 for PEPT1 and 7 for ABHD13), but it has the advantage of increasing the reliability of the method by decreasing the number of templates with a percentage identity of less than 20%.

For ABHD13, 7 templates were used to model 8 strands and 7 helices using 94 alignments (15 multi-template alignments), of which 23 were in the midnight zone (<10%ID), 38 were extreme remote homologues (10<%ID<20), 29 were remote homologues (20<%ID<35) and in only 4 cases was the percentage identity (%ID) greater than 35%. For ABHD16A, 56 unique templates were used to model 9 strands and 15 helices, using 120 alignments (23 multi-template alignments) of which 36 were in the midnight zone, 47 were extreme remote homologues, 19 were remote homologues and in only 10 cases was the percentage identity greater than 35%. Using conventional alignment methods, these alignments would be impossible, but the beauty of the implementation of scaling between 0 and 1 within the multi-template alignment procedure is that small scores occurring at the same position are greatly reduced and high scores are re-enforced. For this reason, it is not necessary for the true alignment to obtain the highest score, provided it obtains a reasonable score within a contributing single-template alignment. In Tables S3 and S4, the percentage identity is shown in **bold** if the alignment score is greater than 0.9 (range 0-1) and in *italics* if it is greater than 025. The first case is indicative of situations where the single template alignments can indicate the correct alignment; the latter is indicative of cases where multi-template alignment can indicate the correct alignment.

The single template method works well into the lower region of the extreme remote homologue zone, as shown in Figure 7, but the use of multiple templates in the alignment extends this considerably because it only requires each template to give the correct alignment a reasonable score rather than the top score, as shown in Figures 5, S5, and S6. The alignments are validated at one of four levels (Table S2), with each level showing how the query structure is compatible with the structural alignment. For ABHD13 and ABHD16A, we have also determined the RMSF using classical all-atom molecular dynamics simulations; this agrees with the positive validation, indicating that the structure is stable.

As shown in Figure 1, some regions of the AI-derived query protein align with many other proteins, while some regions will align with fewer, or possibly no other proteins. The more proteins that align, the greater the options for determining valid templates, but for most ABHD13 and ABHD16A structures, validation was not limited by a lack of suitable templates. If the unaligned regions predominate, it may be pertinent to question whether the fold is genuine.

In conclusion, our new strategy for validating the AI-derived structures of ABHD13 and ABHD16A increased confidence in the MD-based structural interpretations of ABHD13 and ABHD16A in regulating SNAP25 (Mejuto et al., bioRxiv, 2026, doi: 10.64898/2026.01.21.700842).

## References

1 Jumper, J. et al. Highly accurate protein structure prediction with AlphaFold. Nature, doi:10.1038/s41586-021-03819-2 (2021).

2 Tunyasuvunakool, K. et al. Highly accurate protein structure prediction for the human proteome. Nature: p. 596, 590–596, doi:10.1038/s41586-021-03828-1 (2021).

3 Anfinsen, C. B., Redfield, R. R., Choate, W. L., Page, J. & Carroll, W. R. Studies on the gross structure, cross-linkages, and terminal sequences in ribonuclease. J.Biol.Chem.: p. 207, 201–210 (1954).

4 Alexander, L. T. et al. Target highlights in CASP14: Analysis of models by structure providers. Proteins: p. 89, 1647–1672, doi:10.1002/prot.26247 (2021).

5 Buller, R., Damborsky, J., Hilvert, D. & Bornscheuer, U. T. Structure Prediction and Computational Protein Design for Efficient Biocatalysts and Bioactive Proteins. Angew. Chem. Int. Ed. Engl. 64, e202421686, doi:10.1002/anie.202421686 (2025).

6 Graham, F. Daily briefing: AlphaFold developers share Nobel Prize in Chemistry. Nature, doi:10.1038/d41586-024-03325-1 (2024).

7 Callaway, E. Chemistry Nobel goes to developers of AlphaFold AI that predicts protein structures. Nature: p. 634, 525–526, doi:10.1038/d41586-024-03214-7 (2024).

8 Baek, M. et al. Accurate prediction of protein structures and interactions using a three-track neural network. Science: p. 373, 871–876, doi:10.1126/science.abj8754 (2021).

9 Lin, Z. et al. Evolutionary-scale prediction of atomic-level protein structure with a language model. Science: p. 379, 1123–1130, doi:10.1126/science.ade2574 (2023).

10 Zhang, Z., Ou, C., Cho, Y., Akiyama, Y. & Ovchinnikov, S. Artificial intelligencemethods for protein folding and design. Curr. Opin. Struct. Biol. 93, 103066, doi:10.1016/j.sbi.2025.103066 (2025).

11 Abbass, J. From CASP13 to the Nobel Prize: DeepMind’s AlphaFold Journey in Revolutionizing Protein Structure Prediction and Beyond. Curr Protein Pept Sci, doi:10.2174/0113892037374986250711152300 (2025).

12 Webb, B. & Sali, A. Comparative Protein Structure Modeling Using MODELLER. Current protocols in protein science 86, 2.9.1-2.9.37, doi:10.1002/cpps.20 (2016).

13 Eswar, N. et al. Comparative Protein Structure Modeling with MODELLER. Current protocols in bioinformatics, 2.9.1-2.9.31 (2007).

14 Altschul, S. F. et al. Gapped BLAST and PSI-BLAST: a new generation of protein database search programs. Nucleic Acids Res.: p. 25, 3389–3402 (1997).

15 Sievers, F. et al. Fast, scalable generation of high-quality protein multiple sequence alignments using Clustal Omega. Mol. Syst. Biol. 7, Artn 539, doi:10.1038/Msb.2011.75 (2011).

16 Xiang, Z. Advances in homology protein structure modeling. Curr Protein Pept Sci: p. 7, 217–227, doi:10.2174/138920306777452312 (2006).

17 Forrest, L. R., Tang, C. L. & Honig, B. On the accuracy of homology modeling and sequence alignment methods applied to membrane proteins. Biophys. J.: p. 91, 508–517 (2006).

18 Mobarec, J. C., Sanchez, R. & Filizola, M. Modern Homology Modeling of G-Protein Coupled Receptors: Which Structural Template to Use? J. Med. Chem.: p. 52, 5207–5216, doi:10.1021/jm9005252 (2009).

19 Bordin, N. et al. AlphaFold2 reveals commonalities and novelties in protein structure space for 21 model organisms. Communications biology 6, 160, doi:10.1038/s42003-023-04488-9 (2023).

20 Stebbings, L. A. & Mizuguchi, K. HOMSTRAD: recent developments of the Homologous Protein Structure Alignment Database. Nucleic Acids Res.: p. 32 Database issue, D203–D207 (2004).

21 Holm, L. Using Dali for Protein Structure Comparison. Methods Mol. Biol.: p. 2112, 29–42, doi:10.1007/978-1-0716-0270-6_3 (2020).

22 Holm, L. DALI and the persistence of protein shape. Protein Sci.: p. 29, 128–140, doi:10.1002/pro.3749 (2020).

23 Andreeva, A. et al. SCOP database in 2004: refinements integrate structure and sequence family data. Nucleic Acids Res. 32, D226–D229 (2004).

24 Pearl, F. M. et al. A rapid classification protocol for the CATH Domain Database to support structural genomics. Nucleic Acids Res.: p. 29, 223–227 (2001).

25 Hadley, C. & Jones, D. T. A systematic comparison of protein structure classifications: SCOP, CATH and FSSP. Structure Fold.Des: p. 7, 1099–1112 (1999).

26 Orengo, C. A. et al. CATH--a hierarchic classification of protein domain structures. Structure: p. 5, 1093–1108 (1997).

27 Taddese, B. et al. Do plants contain G protein-coupled receptors? Plant Physiol.: p. 164, 287–307, doi:10.1021/ja035737d (2014).

28 Lock, A. et al. One motif to bind them: A small-XXX-small motif affects transmembrane domain 1 oligomerization, function, localization, and cross-talk between two yeast GPCRs. Biochim.Biophys.Acta: p. 1838, 3036–3051 (2014).

29 Reeder, B. J. et al. The circularly permuted globin domain of androglobin exhibits atypical heme stabilization and nitric oxide interaction. Chem. Sci.: p. 15, 6738–6751, doi:10.1039/d4sc00953c (2024).

30 Rost, B. Twilight zone of protein sequence alignments. Protein Eng.: p. 12, 85–94, doi:10.1093/protein/12.2.85 (1999).

31 Hoogewijs, D. et al. Androglobin: A Chimeric Globin in Metazoans That Is Preferentially Expressed in Mammalian Testes. Mol. Biol. Evol.: p. 29, 1105–1114, doi:10.1093/molbev/msr246 (2012).

32 Bracke, A., Hoogewijs, D. & Dewilde, S. Exploring three different expression systems for recombinant expression of globins: Escherichia coli, Pichia pastoris and Spodoptera frugiperda. Anal. Biochem.: p. 543, 62–70, doi:10.1016/j.ab.2017.11.027 (2018).

33 Killer, M., Wald, J., Pieprzyk, J., Marlovits, T. C. & Low, C. Structural snapshots of human PepT1 and PepT2 reveal mechanistic insights into substrate and drug transport across epithelial membranes. Science advances 7, eabk3259, doi:10.1126/sciadv.abk3259 (2021).

34 Kamat, S. S. et al. Immunomodulatory lysophosphatidylserines are regulated by ABHD16A and ABHD12 interplay. Nat. Chem. Biol.: p. 11, 164–171, doi:10.1038/nchembio.1721 (2015).

35 Petropavlovskiy, A. A., Kogut, J. A., Leekha, A., Townsend, C. A. & Sanders, S. S. A sticky situation: regulation and function of protein palmitoylation with a spotlight on the axon and axon initial segment. Neuronal signaling 5, NS20210005, doi:10.1042/NS20210005 (2021).

36 Sanders, S. S. et al. Curation of the Mammalian Palmitoylome Indicates a Pivotal Role for Palmitoylation in Diseases and Disorders of the Nervous System and Cancers. PLoS Comput. Biol. 11, e1004405, doi:10.1371/journal.pcbi.1004405 (2015).

37 Samudrala, R. & Levitt, M. Decoys ‘R’ Us: a database of incorrect conformations to improve protein structure prediction. Protein Sci.: p. 9, 1399–1401, doi:10.1110/ps.9.7.1399 (2000).

38 Wlodarczyk, J. et al. Altered Protein Palmitoylation as Disease Mechanism in Neurodegenerative Disorders. J. Neurosci. 44, doi:10.1523/JNEUROSCI.1225-24.2024 (2024).

39 Lemire, G. et al. ABHD16A deficiency causes a complicated form of hereditary spastic paraplegia associated with intellectual disability and cerebral anomalies. Am. J. Hum. Genet.: p. 108, 2017–2023, doi:10.1016/j.ajhg.2021.09.005 (2021).

40 Zhang, Y. & Skolnick, J. TM-align: a protein structure alignment algorithm based on the TM-score. Nucleic Acids Res.: p. 33, 2302–2309 (2005).

41 Johnson, A. et al. The collaborative UK ECMO trial: Follow-up to 1 year of age. Pediatrics: p. 101 (1998).

