## Supplementary material for "AI-derived Protein Structures Validation: AlphaFold2 Models in the Twilight Zone": Figure S

### Supplementary Information

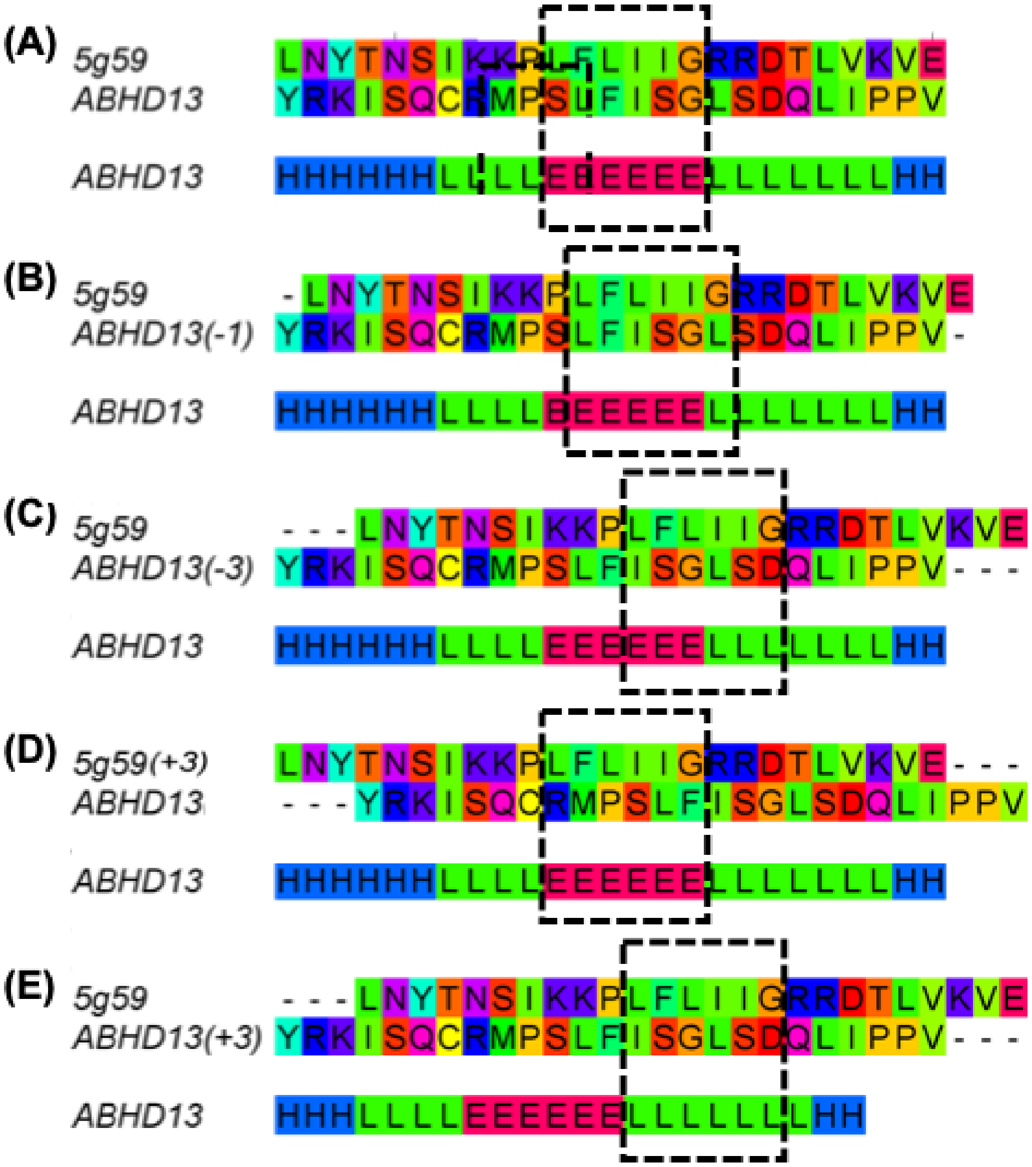


**Figure S1**. Selected pair-wise protein sequence alignments to illustrate the basic Pairwise multi-template local sequence alignments method

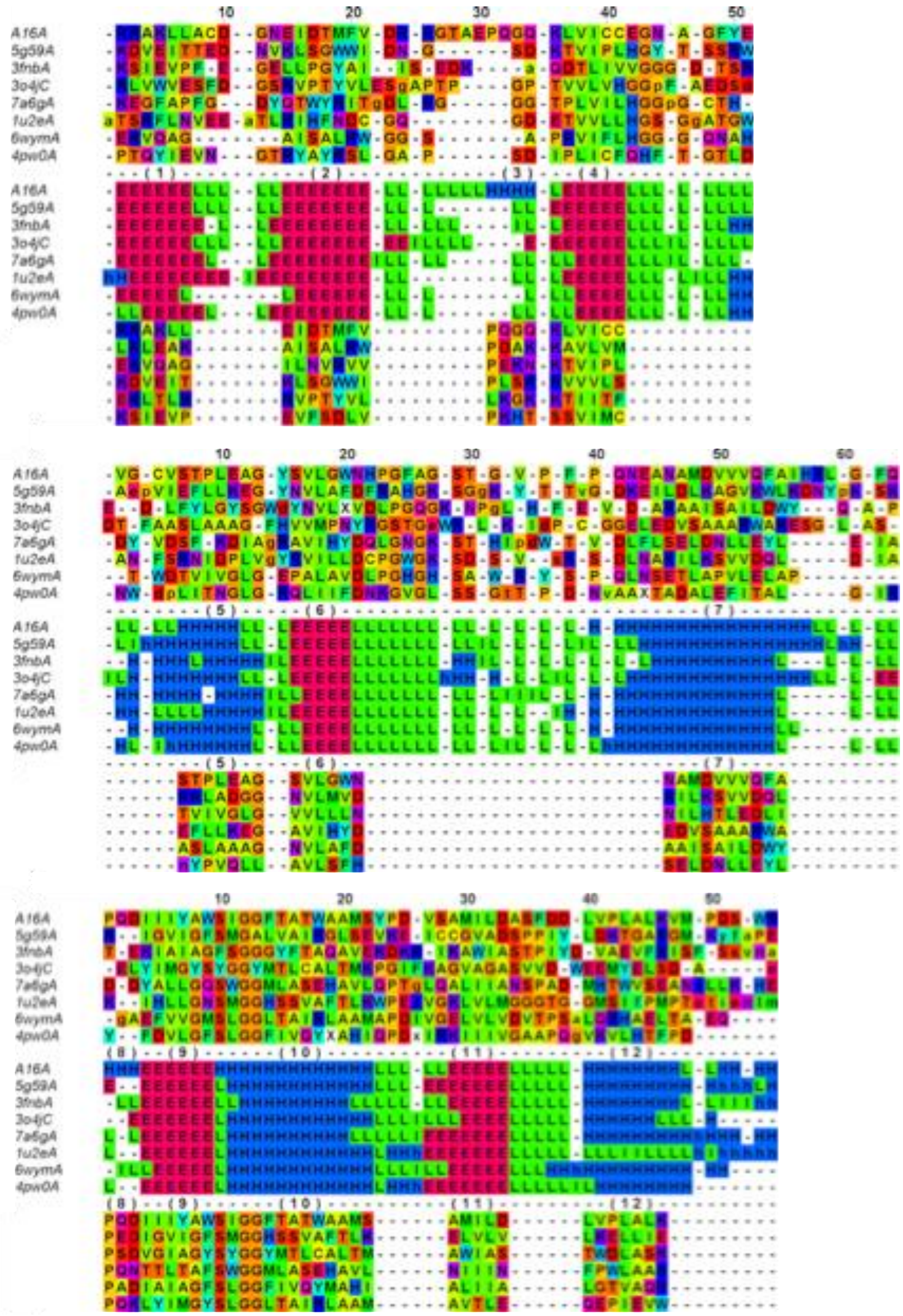

**Figure S2**. The structural alignment between ABHD16A and the 56 templates, as determined by the Dali protein structure comparison server. The templates were chosen because of a relatively high similarity score between the helices and sheets, as determined using the BLOSUM62 similarity matrix.

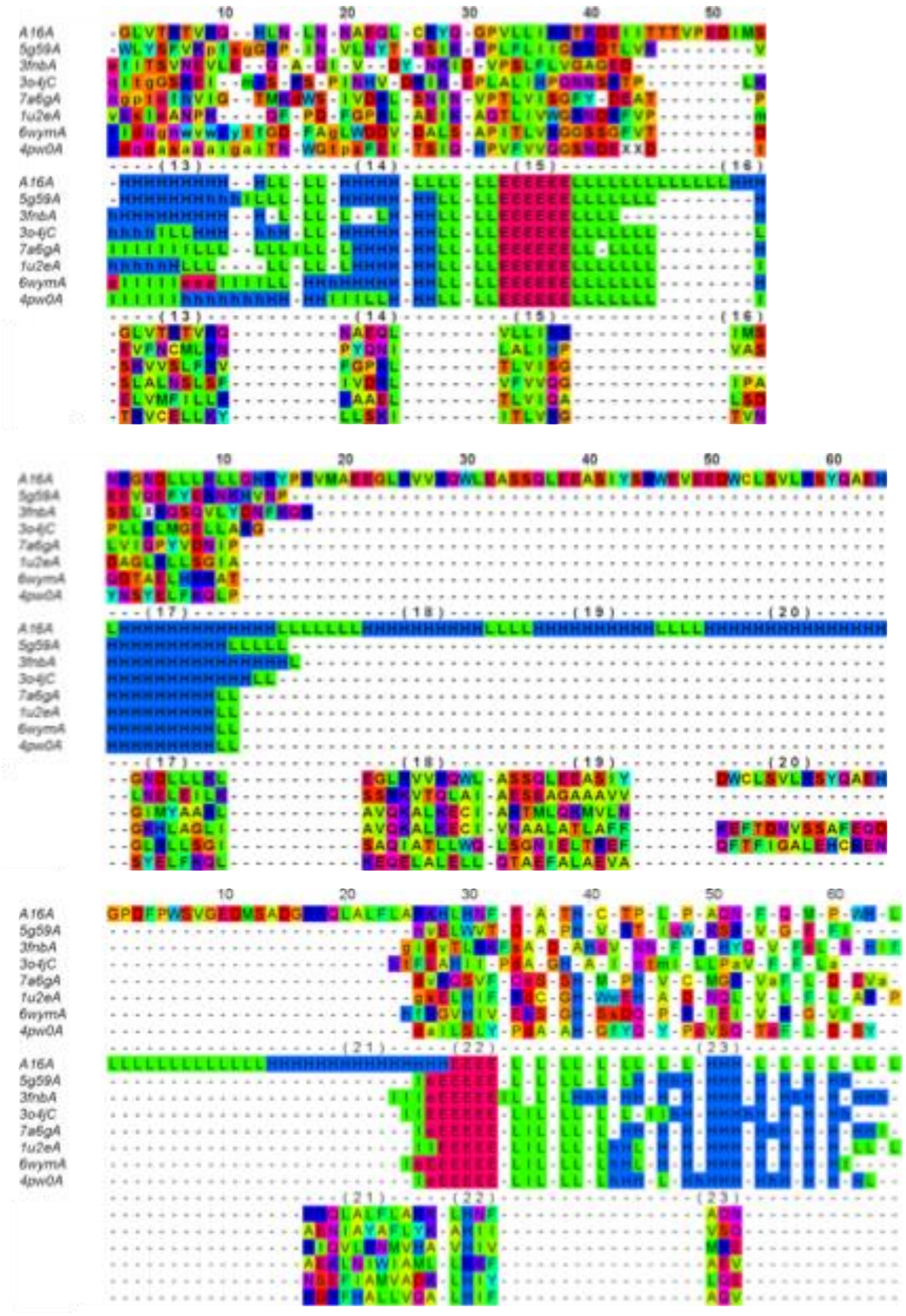


**Figure S3**. The structural alignment between ABHD16A and the 56 templates (continued), as determined by the Dali protein structure comparison server. The templates were chosen because of a relatively high similarity score between the helices and sheets, as determined using the BLOSUM62 similarity matrix.

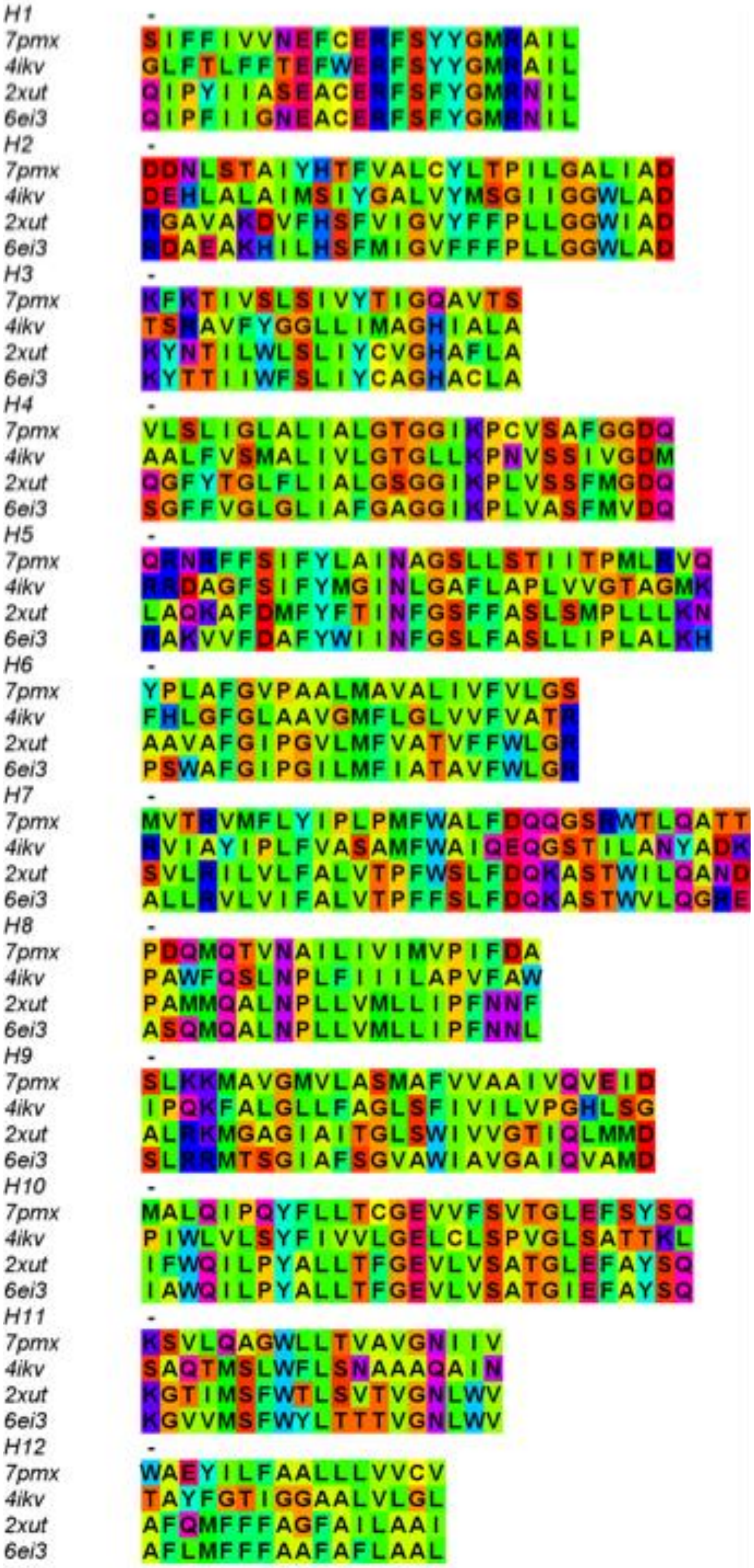

**Figure S4.** Alignment of the 12 transmembrane helices of PEPT1 (pdb code 7pmw) against the 3 templates (pdb codes 4ikv, 2xut, 6ei3).

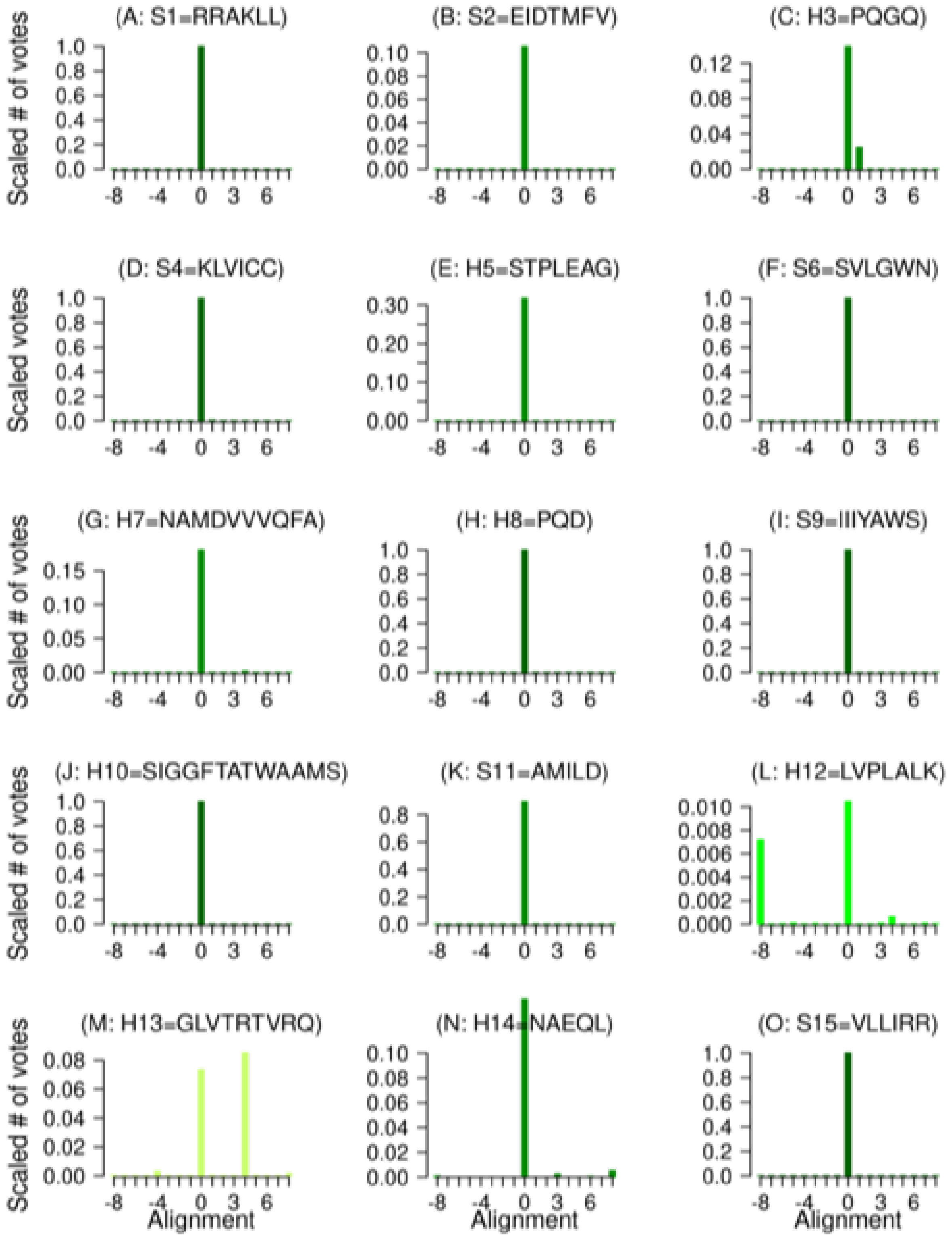


**Figure S5.** Validation of the AlphaFold2 ABHD16 structure, as determined by the multiple template alignment method over the 8 sheets and 15 helical regions. Level 1 validation is shown in dark green (e.g. S1), level 2 in a less dark green (e.g. S2), level 3 in green (e.g. H12), and level 4 validation in dark olive green (e.g. H13). Different templates were identified for each helix or strand due to the low global percentage identity between ABHD16A and the possible templates.

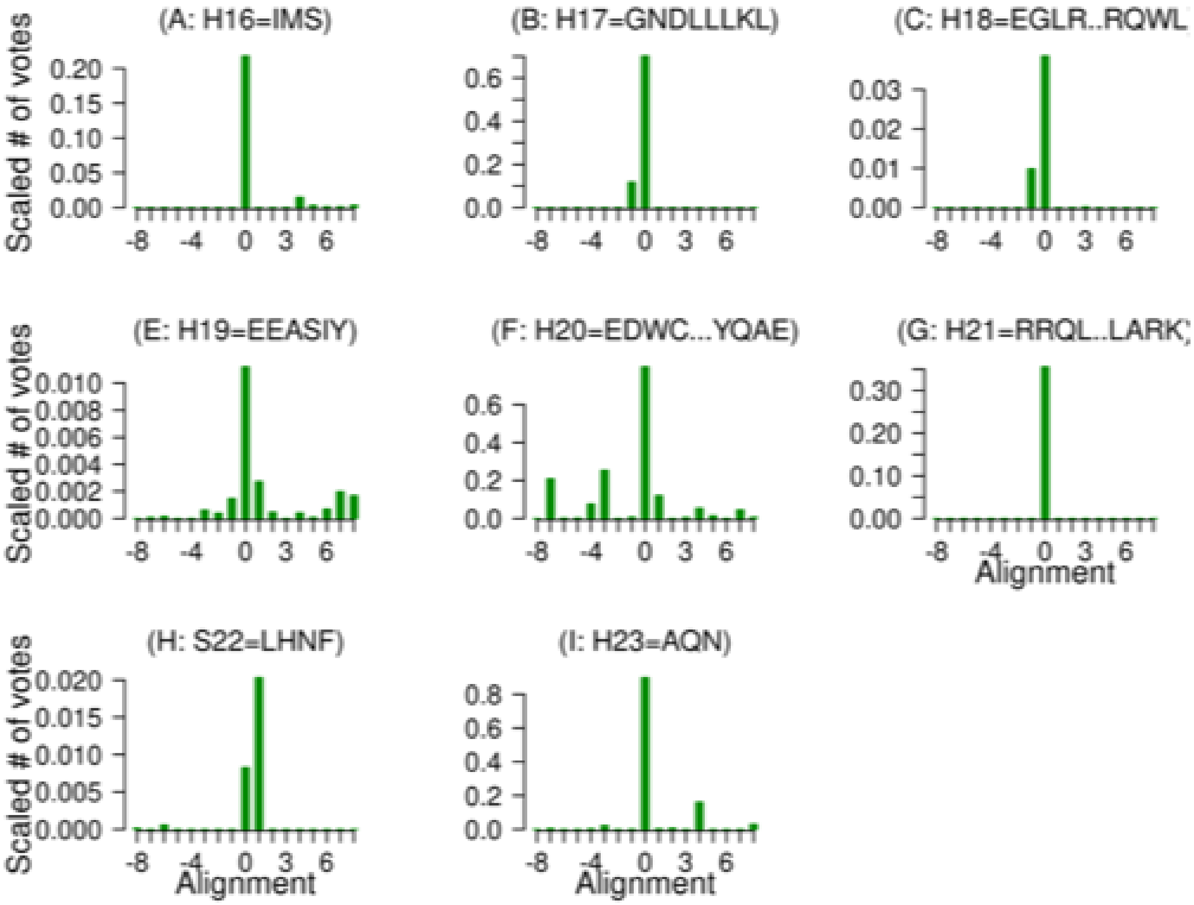


**Figure S6.** Validation of the AlphaFold2 ABHD16 structure, as determined by the multiple template alignment method over the 8 sheets and 15 helical regions. Level 1 validation is shown in dark green (e.g. S1), level 2 in a less dark green (e.g. S2), level 3 in green (e.g. H12), and level 4 validation in dark olive green (e.g. H13). Different templates were identified for each helix or strand due to the low global percentage identity between ABHD16A and the possible templates

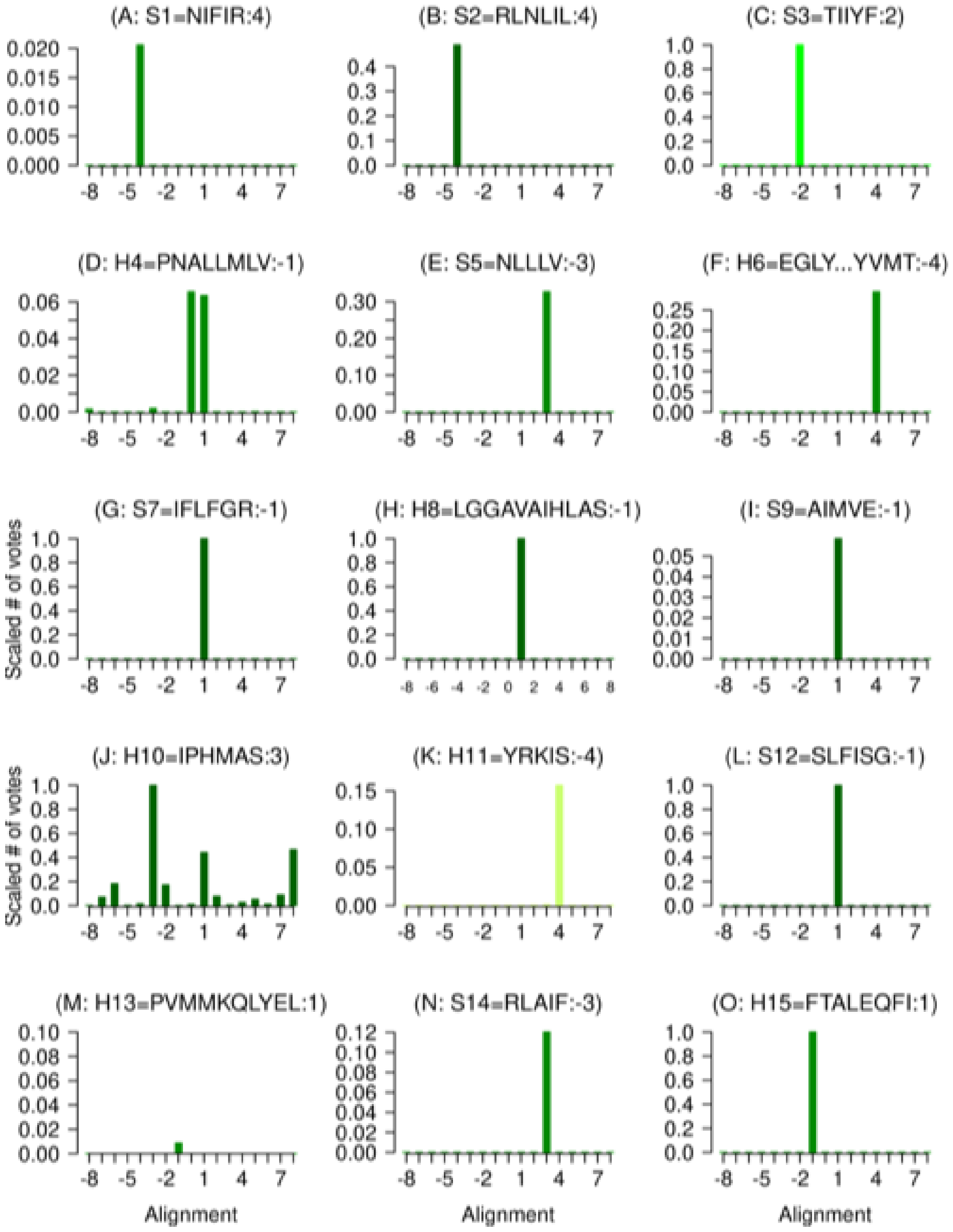


**Figure S7**. Validation strategy applied to a model ABHD13 structure with errors built in to the structural alignment of the helices and sheets. The colour-coded helices and sheets are as in Figure 5; the structural alignment error is indicated in each panel, thus (I: S9=AIMVE:-1) indicates that an alignment error of -1 has been introduced to sheet 9 so the ABHD13 sequence considered is IMVEN

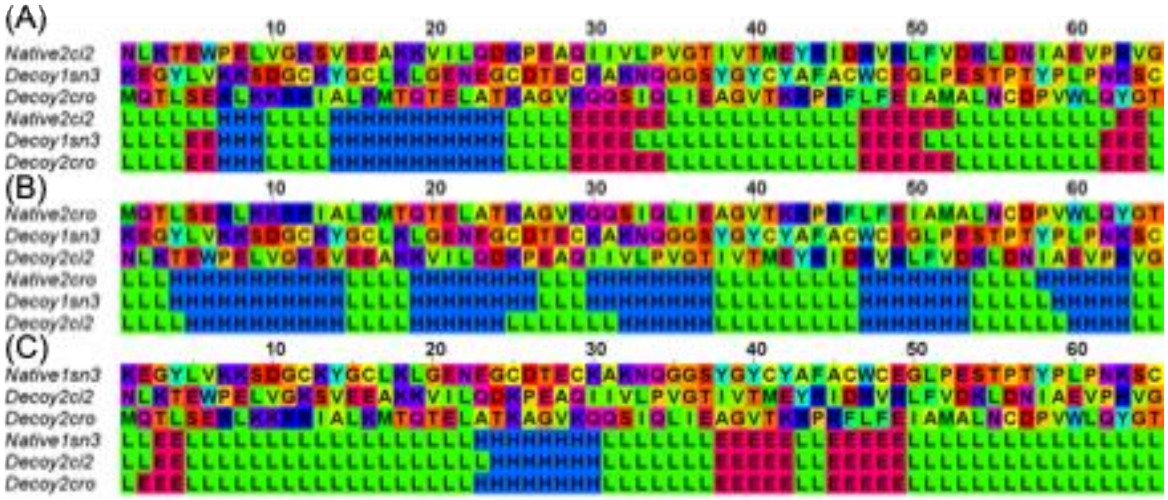


**Figure S8**. Sequence and structural alignment and secondary structure of the six decoy proteins.

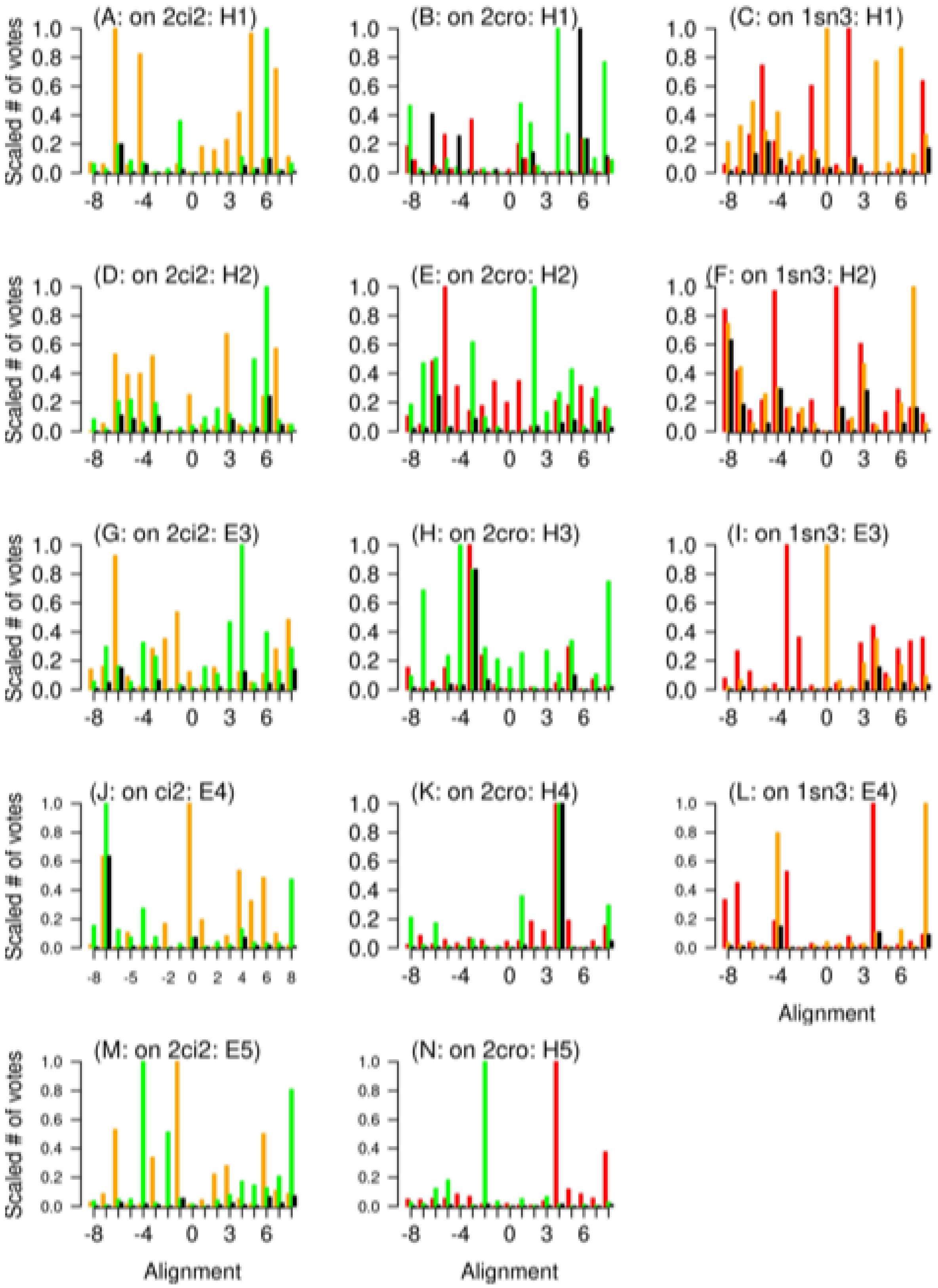


**Figure S9**. Validation strategy applied to misfolded protein models. Left-hand column: the sequence of 2cro and 1sn3 misfolded onto the 2ci2 (red) structure. Middle column: the sequence of 2ci2 and 1sn3 misfolded onto the 2cro (orange) structure. Right-hand column, the sequence of 2cro and 1sn3 (green) misfolded onto the 1sn3 structure. Multiplying the scores for the two template structures gives rise to the black bars.

**Table S1.** BLASTp hits for a search of the Protein Data Bank for the ABHD13 sequence.

| PDB code | E-value | % identity | % coverage |
|---|---|---|---|
| 6ii2A | 4E-09 | 28.3 | 32 |
| 6impA | 6E-09 | 28.3 | 32 |
| 3hjuA | 6E-06 | 28.9 | 48 |
| 7prmA | 7E-06 | 24.4 | 69 |
| 7l4tA | 1E-05 | 24.3 | 69 |
| 3pe6A | 2E-05 | 23.7 | 69 |
| 5zunA | 2E-05 | 23.3 | 76 |
| 3jweA | 3E-05 | 28.3 | 48 |
| 6ax1A | 4E-05 | 28.6 | 48 |
| 3jw8A | 0.001 | 23.1 | 65 |
| 7qjmA | 0.020 | 33.3 | 21 |

**Table S2.** Levels of validation and the associated features. The Table also shows the characteristics associated with misalignment and misfolding.

| Validation Level | Features | Validation Examples from Figures | | |
|---|---|---|---|---|
| | | PEPT1 | ABHD16A | ABHD13 |
| 1 | A single peak at 0 of height close to 1.0 | 4A | 5B | S2A |
| 2 | A single peak at 0 of height less that ~0.9 | N/A | 5F | S2E |
| 3 | 2 or more peaks, where the main peak is at 0 | 4I | 5C | S2L |
| 4 | 2 or more peaks, with a subsidiary peak at 0 with strong representation for 0 peak in individual alignments | N/A | 5K | S2M |
| Misalignment Error | Main peak not at 0. Position of peak indicates extent of misalignment | N/A | S7A | N/A |
| | | Validation on Decoys | | |
| | | 2ci2 | 2cro | 1sn3 |
| Misfolding Error | Main peak not at 0; tendency for multiple strong peaks | S9M | S9B | S9C |

**Table S3. ABHD16A Percentage identities**. **Bold** indicates that the score for the zero alignment is greater than 0.9; *Italics* indicates that the score for the zero alignment is greater than 0.25. The mean percentage identity is 16.1±9.9 across all helices and strands of the 56 templates. The first 4 letters of the motif are shown.

| Temp-<br>late | S1<br>RRAK<br>%ID | Temp-<br>late | S2<br>EIDT<br>%ID | Temp<br>-late | H3<br>PQGQ<br>%ID | Temp-<br>late | S4<br>KLVI<br>%ID |
|---|---|---|---|---|---|---|---|
| 7obm | **23.3** | 6wym | *6.2* | 6kbh | **10.5** | 1jh8 | **15.7** |
| 6wym | **15.7** | 6brt | **4.3** | 6nsi | *10.4* | 5g59 | **24.4** |
| 5g59 | **15.5** | 5g59 | **8.6** | 6qj6 | *8.1* | 6brt | **15.9** |
| 5txc | **16.4** | 3o4j | **11.4** | 4ru0 | **16.8** | 6yuv | **22.0** |
| 3fnb | **10.9** | 5txc | 10.0 | 1zhh | *8.5* | 7dwc | **24.9** |
| Temp-<br>late | H5<br>STPL<br>%ID | Temp-<br>late | S6<br>SVLG<br>%ID | Temp-<br>late | H7<br>NAMD<br>%ID | Temp-<br>late | H8<br>PQD<br>%ID |
| 1q0r | **12.9** | 3fnb | **35.3** | 1u2e | **8.8** | 1dbf | **15.5** |
| 3o4j | **17.0** | 5jrk | **37.2** | 5txc | *7.5* | 1t8h | **13.5** |
| 5g59 | **15.9** | 7a6g | **17.6** | 3o4j | *8.6* | 5thm | **21.8** |
| 4bwg | *10.4* | 5g59 | **20.9** | 3fnb | **17.2** | 5dwd | **21.6** |
| 5jrk | **17.8** | 5txc | **31.9** | 7a6g | **11.0** | 7bft | **20.4** |
| Temp-<br>late | S9<br>IIIY<br>%ID | Temp-<br>late | H10<br>SIGG<br>%ID | Temp-<br>late | S11<br>AMIL<br>%ID | Temp-<br>late | H12<br>LVPL<br>%ID |

| | | | | | | | |
|---|---|---|---|---|---|---|---|
| 5g59 | **41.6** | 1u2e | **38.2** | 6wym | *29.1* | 1vkh | **20.6** |
| 5jrk | **33.3** | 3o4j | **44.8** | 3fnb | **23.9** | 7oqe | 5.8 |
| 7obm | **34.0** | 7a6g | **37.0** | 6yuv | **12.1** | 2yys | *9.6* |
| 3fnb | **39.5** | 4pw0 | **40.9** | 7a6g | **31.8** | 6oyc | **20.3** |
| 3o4j | **35.4** | 6wym | **39.4** | 7obm | **18.5** | 5g59 | *8.0* |
| Template | H13 GLVT %ID | Template | H14 NAEQ %ID | Template | S15 VLLI %ID | Template | H16 IMS %ID |
| 4j8s | **9.0** | 7obm | *8.5* | 3o4j | **17.2** | 1vhd | **16.8** |
| 5w66 | *9.4* | 1u2e | **15.9** | 7a6g | **20.3** | 5fb3 | **12.5** |
| 2a3v | 6.4 | 7a6g | **18.9** | 4pw0 | **18.8** | 2q01 | **12.5** |
| 5yu7 | **18.2** | 1q0r | **16.4** | 1q0r | **16.5** | 1thg | *6.6* |
| 4x4w | *7.9* | 6yuv | **12.3** | 6wym | **27.9** | 5zl9 | *8.1* |
| Template | H17 GNDL %ID | Template | H18 EGLR %ID | Template | H19 EEAS %ID | Template | H20 DWCL %ID |
| 6yuv | **9.6** | 5dcq | *7.9* | 1k5d | *7.7* | 8ou0 | **9.2** |
| 7dwc | *13.6* | 3cfo | *11.2* | 1gvn | *5.7* | 3bvo | *9.7* |
| 1q0r | **16.0** | 3bvo | 8.1 | 2oeb | **9.6** | | |
| 1u2e | **14.7** | 6rdj | **9.4** | 3sae | 5.4 | | |
| 4pw0 | **15.8** | 2k9m | *8.4* | 6fht | 7.0 | | |
| Template | H21 RRQL %ID | Template | S22 LHNF %ID | Template | H23 AQN %ID | | |
| 2oeb | *11.0* | 3o4j | *9.5* | 4pw0 | **9.4** | | |
| 4wzi | **20.0** | 6wym | *7.6* | 1q0r | **12.1** | | |
| 5nlg | **10.4** | 3fnb | 11.1 | 7dwc | *8.2* | | |
| 1fc3 | *3.4* | 7dwc | *10.4* | 6brt | **10.7** | | |
| 6g7c | **17.1** | 1u2e | *9.5* | 2o2g | **16.5** | | |

**Table S4. ABHD13 Percentage identities**. **Bold** indicates that the score for the zero alignment is greater than 0.9; *Italic* indicates that the score for the zero alignment is greater than 0.25. Helix 7 and Helix 15 were shortened by 2 and 4 residues, respectively, due to gaps in the MSAs. Missing templates, e.g. 2i3d for helix 11, arise due to gaps in the appropriate region of the sequence alignment. The average percentage identity is 16.6$\pm$9.0 across all the helices and strands of the 7 templates; this is well within the twilight zone. The minimum percentage identity is 5.4 for helix 11 of 5hdf; the maximum percentage identity is 27.6 for strand 12 of 2o2g 2o2g.

| | Sht 1 | Sht 2 | Sht 3 | Hlx 4 | Sht 5 | Hlx 6 | Sht 7 | Hlx 8 |
|---|---|---|---|---|---|---|---|---|
| Protein | NIFIR | RLNLIL | TIIYF | PNALLMLV | NLLLV | EGLYLDSEA VLDYVM | IFLFGR | LGGAV AIHLAS |
| 2i3d | **16.9** | **18.0** | *10.3* | **12.6** | **12.3** | **27.5** | **25.0** | **24.7** |
| 2o2g | **16.2** | **23.9** | *18.4* | **14.0** | **13.6** | **12.1** | **31.0** | **32.2** |
| 3pf8 | **10.8** | **24.2** | **17.3** | **13.0** | *15.0* | **18.9** | **27.4** | **36.5** |
| 4ao8 | **14.1** | **13.7** | *20.1* | **12.7** | **16.0** | **19.4** | **16.0** | **24.7** |
| 4xoo | **9.5** | **14.4** | **18.4** | **8.1** | **13.2** | **15.2** | **25.0** | **32.2** |
| 5g59 | **21.0** | **26.6** | **24.8** | - | **31.0** | **20.5** | **32.0** | **32.5** |
| 5hdf | **14.7** | **15.9** | 7.3 | **7.8** | **12.7** | *7.0* | **29.0** | **26.2** |
| Mean | 14.7 | 19.5 | 16.6 | 11.4 | 16.3 | 14.3 | 26.5 | 29.6 |

| | Sht 9 | Hlx 10 | Hlx 11 | Sht 12 | Hlx 13 | Sht 14 | Hlx 15 | Mean |
|---|---|---|---|---|---|---|---|---|
| Protein | AIMVE | IPHMAS | YRKIS | SLFISG | PVMMKQLYEL | RLAIF | FTALEQFI | |
| 2i3d | **21.2** | - | - | **29.9** | **10.7** | 5.5 | **12.0** | 17.4 |
| 2o2g | **16.4** | - | **8.7** | **37.6** | *6.7* | *7.7* | *6.4* | *17.1* |
| 3pf8 | **19.5** | **9.3** | **9.3** | **32.4** | **9.4** | 5.7 | - | 17.7 |
| 4ao8 | **18.3** | - | *6.3* | **21.6** | **16.4** | *7.0* | **16.9** | 15.1 |
| 4x00 | **23.8** | - | **7.6** | **36.0** | - | *6.3* | **18.1** | 17.2 |
| 5g59 | **27.1** | **12.7** | **14.3** | **35.8** | - | 5.9 | - | 22.6 |
| 5hdf | **27.4** | | *5.4* | **28.6** | *5.5* | *7.6* | **14.6** | 15.0 |
| Mean | 20.3 | 10.7 | 8.6 | 31.7 | 9.7 | 6.5 | 13.6 | 17.4 |
